# Deletion of the Envelope gene attenuates SARS-CoV-2 infection by altered Spike localization and increased cell-to-cell transmission

**DOI:** 10.1101/2025.05.20.655126

**Authors:** Hannah L. Fischer, Shanmugapriya Shanmugapriya, Christopher Kline, W. Paul Duprex, Kevin R. McCarthy, Simon C. Watkins, James F. Conway, Zandrea Ambrose

## Abstract

Severe acute respiratory syndrome coronavirus 2 (SARS-CoV-2) causes COVID-19, a highly transmissible acute respiratory infection that can result in severe pneumonia and death. Many details of SARS-CoV-2 infection are not fully understood, including the cell biology and host-virus interactions involved in coronavirus assembly and release, in which the Envelope (E) structural protein is instrumental. Deletion of E in other coronaviruses has been shown previously to either attenuate or abrogate infection. To determine the role of E on SARS-CoV-2 virus production and infectivity, we produced reporter SARS-CoV-2 with or without the E gene deleted using a bacterial artificial chromosome. Replication of ΔE SARS-CoV-2 was attenuated in Vero E6 cells expressing human ACE2 and TMPRSS2 and in human epithelial cell lines. Electron and immunofluorescence microscopy and virology assays showed that ΔE SARS-CoV-2 increased cell surface expression of Spike (S) glycoprotein, leading to reduced S incorporation into ΔE SARS-CoV-2 particles and promotion of increased cell-to-cell transmission that evades neutralizing antibody inhibition. Trans-complementation of E partially rescued ΔE SARS-CoV-2 S incorporation and restored cell-free transmission. In addition to validating the role of E in retention of S in the ER-Golgi intermediate complex (ERGIC), our results showed that a lack of E led to reorganization of the ERGIC and Golgi during SARS-CoV-2 infection. Improved understanding of E in SARS-CoV-2 replication and host pathogenesis may help development of novel therapeutics.

**Importance:** Non-Spike coronavirus structural proteins, including Envelope, are conserved, making them potential pan-coronavirus therapeutic targets. Many details about these proteins and their roles in viral replication and host pathogenesis are unknown. In this study, we showed that SARS-CoV-2 replicates without Envelope but is attenuated and impaired for virus particle formation, with less Spike incorporated into virions and more Spike expressed on the cell surface compared to wild-type virus. SARS-CoV-2 lacking Envelope transmitted primarily via cell fusion and evaded neutralizing antibodies. In addition, the absence of Envelope resulted in the reorganization of the cell secretory compartment during SARS-CoV-2 infection. A better understanding of how Envelope influences SARS-CoV-2 replication could guide directed design of novel therapeutics for treatment of COVID-19 patients, as well as the potential for pan-coronavirus protection against future coronavirus outbreaks.

## Introduction

Severe acute respiratory syndrome coronavirus type 2 (SARS-CoV-2) is the third deadly coronavirus (CoV) to emerge in humans from zoonotic sources (1), preceded by SARS-CoV-1 in 2002 (2) and Middle Eastern respiratory syndrome coronavirus (MERS-CoV) in 2012 (3). Infection with SARS-CoV-2 causes COVID-19, an acute respiratory infection that presents with mild to moderate symptoms in most individuals but can result in severe pneumonia, multisystem inflammatory syndrome, multiple organ failure, and death (4–9). Since the start of the global pandemic in late 2019, there have been over 779 million cases of COVID-19 and over 7.1 million deaths reported worldwide as of December 2025 (10). Post-acute sequelae are prominent with COVID-19, with over 10% of infected individuals developing Long COVID, consisting of over 200 identified symptoms, including long-term metabolic, cardiovascular, and neurological morbidities (11, 12).

Achievements in preventing and treating COVID-19 were rapidly developed to combat the pandemic. SARS-CoV-2 vaccines have been successful in preventing COVID-19, though protection conferred by either vaccines or natural infection lack long-term durability (13, 14). This is confounded by the rapid development of SARS-CoV-2 variants that evade immunity, requiring development and administration of vaccine boosters that target circulating variants, similar to seasonal influenza viruses (15–17). Most current treatments for COVID-19 in the U.S. are high-titer convalescent plasma or nonspecific small molecule antivirals that consist of repurposed inhibitors targeting viral enzymes common across multiple RNA viruses (18). Improved SARS-CoV-2 therapeutics are needed for higher specificity and efficacy, ideally against conserved CoV proteins that may offer pan-coronaviral protection.

SARS-CoV-2 is a β-coronavirus that is primarily transmitted via respiratory droplets and aerosols (19, 20) , infecting cells expressing the ACE2 receptor, most commonly epithelial cells within the upper airways and the gastrointestinal tract (21–24). The viral Spike (S) glycoprotein binds to ACE2 and is primed by the host protease furin and subsequently activated for fusion and entry primarily by the host protease TMPRSS2 or, alternatively, Cathepsin B/L (25–27). Cytoplasmic release of the 30 kb single-stranded, positive-sense RNA viral genome results in ribosomal translation of the ORF1a/b open reading frame into the pp1a and pp1ab polyproteins, which are cleaved by two virally encoded proteases into the 16 nonstructural proteins (Nsps) comprising the viral replicase machinery (28, 29). Interaction of Nsps with host factors results in the formation of double membrane vesicles, in which the viral genome is both replicated and used as a template for discontinuous transcription of nested, co-terminal subgenomic RNAs (sgRNAs) that are translated into the viral structural and accessory proteins (28, 29). The membrane-bound S, Envelope (E), and Membrane (M) structural proteins along with the Nucleocapsid (N)-coated viral RNA genome are assembled into nascent virions within the endoplasmic reticulum/Golgi intermediate compartment (ERGIC) and released from the cell via vesicles (28, 29).

E is the smallest structural protein with only 75 amino acids, consisting of a short, luminal N-terminal domain (NTD); a transmembrane domain (TMD) that forms a calcium-selective channel in the lipid bilayer of the ERGIC membrane; and a cytosolic C-terminal domain (CTD) containing a PDZ-binding motif (PBM) that enables interaction with PDZ-containing host proteins (30–33). Expression of recombinant SARS-CoV-1 E by transfection or during infection of cells results in colocalization with ERGIC proteins and it is not found in the ER or at the plasma membrane (34). E is conserved across CoVs and has three demonstrated roles in CoV replication: virus assembly, virus release, and host pathogenesis (35–39). However, the details for these roles in CoV infection are not completely understood.

The requirement of E for replication varies significantly between CoVs. Deletion of the E gene from transmissible gastroenteritis virus (TGEV), human coronavirus OC43 (HCoV-OC43), and MERS-CoV prevents viral propagation (40–42), whereas deletion of E in murine hepatitis virus (MHV) and SARS-CoV-1 only attenuate infection via an unknown mechanism (43, 44). While deleting the ORF3a and E genes from SARS-CoV-1 and SARS-CoV-2 eliminates viral propagation (45, 46), the effects of deleting only E from SARS-CoV-2 have not been reported.

Here we describe the development of a bacterial artificial chromosome (BAC) encoding SARS-CoV-2 with nanoluciferase (nLuc) in place of ORF7a/b to produce infectious reporter virus. We used this BAC system with the E gene deleted from the viral genome, ΔE SARS-CoV-2, which resulted in virus with attenuated replication. Virus particles produced in the absence of E incorporated less S and were impaired for cell-free transmission. Trans-complementation of E partially rescued S incorporation and infection via virus particle spread. The primary mechanism of ΔE SARS-CoV-2 transmission was through cell fusion, which permitted evasion of antibody neutralization. ΔE SARS-CoV-2 infection resulted in greater cell surface expression of S and reorganization of the ERGIC and Golgi.

## Results

### Design and construction of a BAC-based SARS-CoV-2 virus production system

We designed a mammalian-expression BAC encoding the complete USA-WA1/2020 SARS-CoV-2 genome (**Fig S1A**), similar to the method reported with SARS-CoV-1 and MERS-CoV (42, 47). The ORF7a/b gene was replaced with the codon-optimized nLuc reporter gene (SARS-CoV-2-nLuc) in the same manner previously reported with a SARS-CoV-2 mNeonGreen reporter virus (48) and a SARS-CoV-1 GFP reporter virus (49). The CopyControl™ pCC1BAC vector was chosen due to its dual origin of replication that optimizes insert stability under normal bacteria propagation conditions but also permits higher DNA plasmid yield under chemical induction (50). As pCC1BAC is not designed for mammalian expression, we synthesized a mammalian expression cassette (MEC) containing the necessary *cis*-acting elements to regulate expression of the SARS-CoV-2 genome upon transfection of the BAC into mammalian cells, and to alternatively promote *in vitro* RNA synthesis from the BAC for transfection of genomic viral RNA into mammalian cells. Specifically, the human cytomegalovirus (CMV) immediate early promoter and bacteriophage T7 RNA polymerase promoter were engineered upstream of the insert site and a polyA(29) tail, the hepatitis D virus (HDV) antigenomic ribozyme, and bovine growth hormone (bGH) polyadenylation signal were engineered downstream of the insert site. Herein, WT SARS-CoV-2 virus refers to this recombinant USA-WA1/2020, while WT SARS-CoV-2-nLuc refers to recombinant USA-WA1/2020 in which ORF7a/b was replaced with nLuc.

### SARS-CoV-2 BAC system produces infectious virus

Production of SARS-CoV-2 and SARS-CoV-2-nLuc was performed by transfection of the BAC into BHK cells, which are highly transfectable but not permissive to SARS-CoV-2 infection (**Fig S1B**). The transfected BHK cells were overlayed onto Vero E6 cells stably expressing human ACE2 and TMPRSS2 (Vero E6-TMPRSS2-T2A-ACE2 cells, referred to herein as Vero E6-hAT cells) to amplify virus production. SARS-CoV-2-nLuc was also produced by using the BAC as the template for *in vitro* RNA synthesis and transfecting the full-length, capped viral RNA into BHK cells (data not shown).

Vero E6-hAT cells infected with SARS-CoV-2-nLuc produced from the BAC exhibited cytopathic effects (CPE), viral plaques, and an infectious titer typically observed with SARS-CoV-2 infection (**Fig 1A-B**). Genomic RNA copies were detected in both infected cells and viral supernatant (**Fig 1C-D**). Comparable results were observed with SARS-CoV-2 virus lacking the nLuc reporter gene (i.e., with the ORF7a/b gene intact) (**Fig S2A-D**). Similarly, nLuc expression was detected in infected cells (**Fig 1E**). In addition, RNA was isolated from infected Vero E6-hAT cells and amplified by RT-PCR using a forward primer specific to the common 5’-UTR leader sequence and reverse primers specific to each sgRNA. Agarose gel electrophoresis demonstrated that all sgRNAs were produced in cells infected with SARS-CoV-2-nLuc (**Fig 1F**). Negative stain electron microscopy (EM) and cryogenic EM (Cryo-EM) of SARS-CoV-2-nLuc supernatant showed prototypical coronavirus particles of approximately 125 nm diameter and decorated with S glycoprotein trimers (**Fig 1G**). Altogether, these results demonstrate that the SARS-CoV-2 BAC system produces infectious virus particles that replicate in permissive cells.

**Figure 1.**
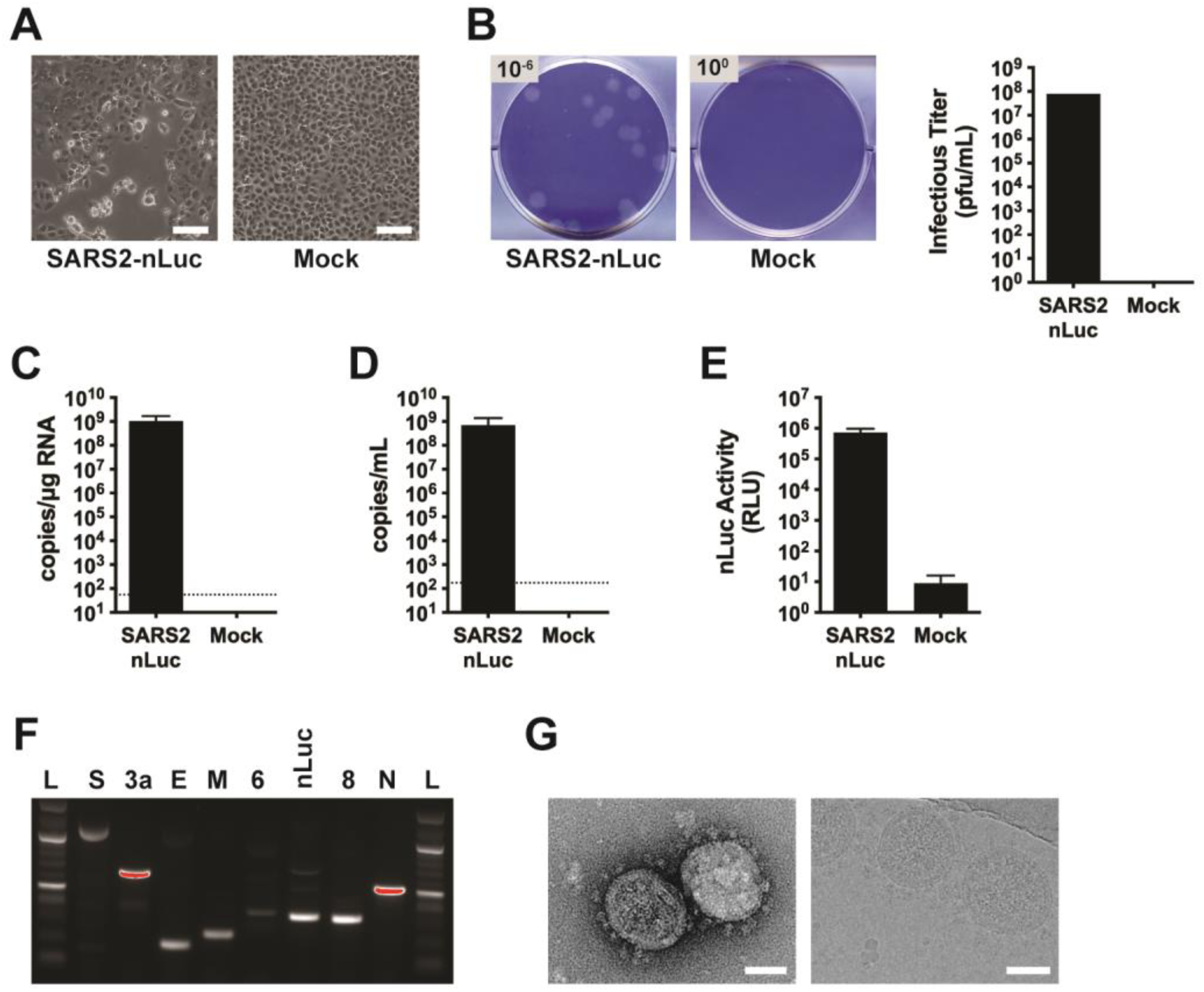
SARS-CoV-2 BAC system produces infectious virus particles. (A) Vero E6-hAT cells were infected with WT SARS-CoV-2-nLuc virus (MOI=1) or mock infected for 24 h and imaged by phase contrast microscopy with a 10X objective. Solid white bar represents 100 μm. (B) Plaque titer of WT SARS-CoV-2-nLuc virus on Vero E6-hAT cells. (C,D) RNA was isolated from WT SARS-CoV-2-nLuc virus producer cells (C) and supernatant (D) and ORF1a copies were quantified by qRT-PCR. The dotted lines represent the assay limits of detection. Error bars represent standard error of the mean (SEM) for two independent experiments. (E) E6-hAT cells were infected with WT SARS-CoV-2-nLuc virus or mock infected for 48 h and lysates were measured for nLuc activity. Error bars represent SEM for three independent experiments. (F) RNA was isolated from Vero E6-hAT cells infected with WT SARS-CoV-2-nLuc virus and RT-PCR amplified with SARS-CoV-2 sgRNA specific primers and run on an agarose gel. L = 100 bp ladder. (G) Negative stain EM (left) and Cryo-EM (right) of WT SARS-CoV-2-nLuc virus particles. Solid white bar represents 50 nm.

### Construction of ΔE SARS-CoV-2 BAC and E trans-complementation cell lines

To examine the role of E in SARS-CoV-2 infection, the first 186 bp of the E gene was together with the transcription regulatory sequence (TRS) upstream of the E gene were deleted in the SARS-CoV-2 and SARS-CoV-2-nLuc BACs (**Fig S3A**). The remaining 42 bp of E were left intact to preserve expression of the downstream M gene by ensuring the M gene TRS and surrounding RNA structure were unaltered.

To complement the deleted E gene *in trans*, stable BHK and Vero E6-hAT cell lines were engineered to inducibly express E. To minimize the likelihood of recombination of the trans-complemented E mRNA with the ΔE SARS-CoV-2 viral genome, the E gene was codon-optimized (coE), which removed sequence homology. To facilitate identification of cells expressing E, the codon-optimized mRuby3 fluorescent protein was linked to the N-terminus of coE by the 2A self-cleaving peptide from *Thosea asigna* virus (T2A), permitting co-expression of mRuby3 and coE (**Fig S3B**). The mRuby3-T2A-coE construct was cloned into doxycycline-inducible lentiviral vectors and transduced into BHK and Vero E6-hAT cells. Prior to each use, the cell lines were induced with doxycycline for 48h and visually confirmed by microscopy to express mRuby3, and therefore also coE (**Fig S3B**). Expression of coE protein was confirmed by immunoblot (**Fig S3C**).

### ΔE SARS-CoV-2 is replication competent but attenuated compared to wild-type virus

ΔE SARS-CoV-2 and SARS-CoV-2-nLuc lacking E incorporation were produced as described for WT SARS-CoV-2 above. To produce ΔE SARS-CoV-2 trans-complemented with E protein, the ΔE SARS-CoV-2 BAC was transfected into induced BHK-coE cells that were overlaid onto induced Vero E6-hAT-coE cells (**Fig S3D**). A multi-passage workflow was developed to characterize ΔE SARS-CoV-2 infectivity and replication (**Fig S3E**). The initial passages (P1) of WT and ΔE SARS-CoV-2-nLuc viruses (the latter with and without E trans-complementation) were inoculated onto fresh Vero E6-hAT and Vero E6-hAT-coE cells (P2), which were subsequently passaged a third time (P3). For each passage, both the virus supernatant and the producer cells were collected and characterized.

Plaque titration revealed ΔE SARS-CoV-2-nLuc without E trans-complementation produced moderate virus titers (10^4^ to 10^5^ pfu/mL) in Vero E6-hAT cells, which were attenuated 2- to 3-log compared to WT SARS-CoV-2-nLuc (**Fig 2A**). Similarly, quantitative RT-PCR (qRT-PCR) of genomic RNA copies in viral supernatant showed 2- to 4-log lower levels (**Fig 2B**). Comparable results were observed with ΔE SARS-CoV-2 without the nLuc reporter gene (**Fig S2C-D**). Infecting fresh Vero E6-hAT cells with each virus further demonstrated a 3- to 4-log reduction in nLuc activity for ΔE SARS-CoV-2-nLuc compared to WT (**Fig 2C**). However, complementing E *in trans* did not significantly increase infectivity, genomic RNA abundance, or nLuc activity (**Fig 2A-C**).

**Figure 2.**
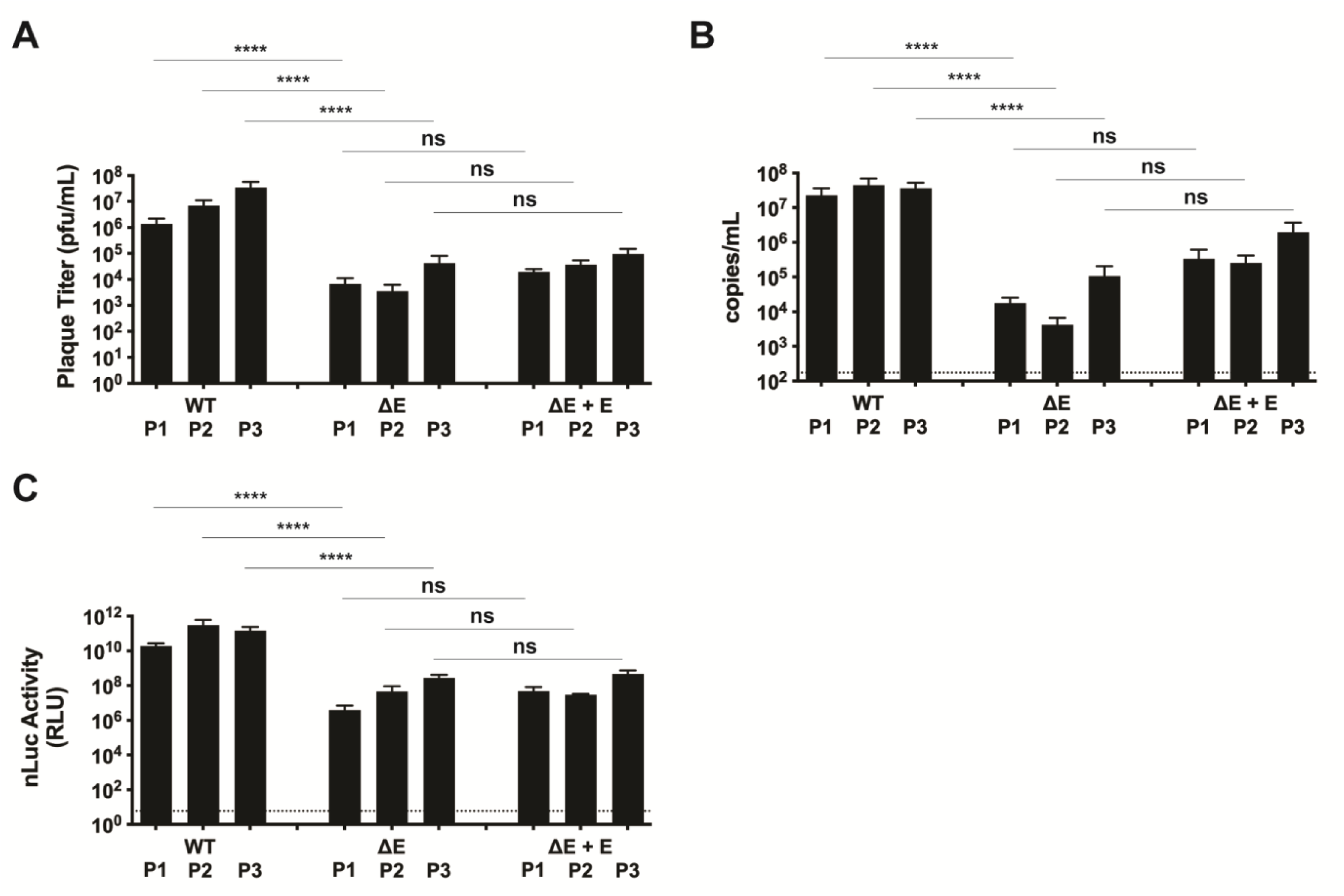
ΔE SARS-CoV-2 is replication-competent but attenuated compared to WT virus. (A-C) WT and ΔE SARS-CoV-2-nLuc viruses were serially passaged three times in Vero E6-hAT cells (WT and ΔE) or E6-hAT cells stably expressing coE *in trans* (ΔE only). Dotted lines represent the assay limits of detection. **** p < 0.0001, ns p > 0.05. Error bars represent SEM for three independent experiments. Plaque titers (A), genomic RNA (ORF1a) copies (B), and infectivity on fresh Vero E6-hAT cells after 72 h (C) were measured for each passage.

As Vero E6-hAT cells artificially express human ACE2 and TMPRSS2, they do not represent a biologically relevant target cell of SARS-CoV-2 infection. To determine whether the observed attenuated phenotype of ΔE SARS-CoV-2 is cell type specific and to examine infection in more relevant cells, human epithelial colorectal Caco-2 cells and human respiratory epithelial Calu-3 and 16HBE cells were infected with WT and ΔE SARS-CoV-2-nLuc and assayed for nLuc activity. A similar 1.5- to 3-log reduction in infectivity was observed for ΔE SARS-CoV-2-nLuc compared to WT in Vero E6, Caco-2, and Calu-3 cells as was observed in Vero E6-hAT cells (**Fig 3A**).

**Figure 3.**
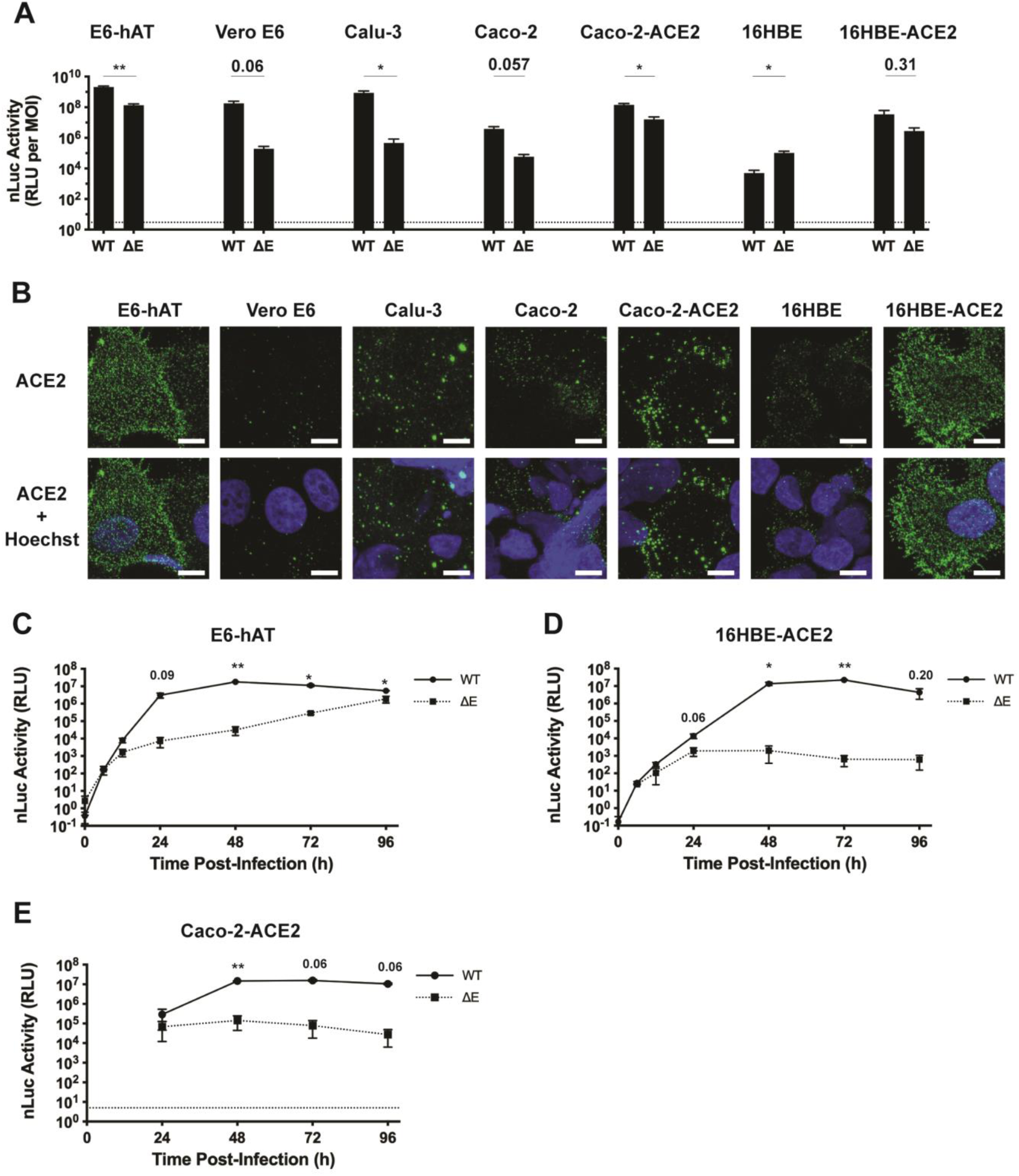
ΔE SARS-CoV-2 replication is attenuated in human epithelial cell lines compared to WT virus. (A) Indicated cell lines were infected with WT or ΔE SARS-CoV-2-nLuc and assayed for nLuc activity after 72 h. Dotted lines represent the assay limits of detection. ** p < 0.01, * p < 0.05. Error bars represent SEM for 2-3 independent experiments. (B) Representative images of indicated cell lines that were fixed, stained for ACE2 cell surface expression (green) and nuclei (Hoechst; blue), and imaged by confocal microscopy. Solid white bar represents 10 μm. Vero E6-hAT cells (F), 16HBE-ACE2 cells (G), or Caco-2-ACE2 cells (H) were infected with WT or ΔE SARS-CoV-2-nLuc and assayed for nLuc activity after indicated time points. ** p < 0.01, * p < 0.05. Error bars represent SEM for 2-3 independent experiments.

Surprisingly, while ΔE SARS-CoV-2-nLuc had similar infectivity in 16HBE cells as in the other cell lines, WT SARS-CoV-2-nLuc infectivity was about 1-log lower than ΔE SARS-CoV-2-nLuc infectivity (**Fig 3A**). To investigate whether 16HBE cells had lower expression of ACE2, all cell lines were stained for human ACE2 and imaged for immunofluorescence (**Fig 3B**). Vero E6-hAT cells, as expected, showed high ACE2 expression, commensurate with the observed infectivity of SARS-CoV-2-nLuc. The human cell lines showed lower levels of ACE2 expression than Vero E6-hAT cells, with 16HBE cells having the lowest ACE2 expression. Parental Vero E6 cells, which had relatively high WT SARS-CoV-2-nLuc infectivity, showed low surface expression of ACE2.

To determine if increased ACE2 expression could rescue WT SARS-CoV-2-nLuc infectivity in 16HBE cells, they were transduced with a lentiviral vector encoding human ACE2, resulting in a similar level of ACE2 cell surface expression as in Vero E6-hAT cells (**Fig 3B**). Both WT and ΔE SARS-CoV-2-nLuc showed increased infectivity in 16HBE-ACE2 cells compared with parental 16HBE cells (**Fig 3A**); however, while ΔE SARS-CoV-2-nLuc infectivity was about 1-log higher in 16HBE-ACE2 cells, WT SARS-CoV-2-nLuc infectivity was enhanced 3.5-log in 16HBE-ACE2 cells. Similar enhancement of ACE2 expression and SARS-CoV-2-nLuc infectivity was observed with Caco-2 cells that were transduced with the human ACE2 lentiviral vector (**Fig 3A-B**). As these results were all obtained at a single, late time point post-infection, Vero E6-hAT, 16HBE-ACE2, and Caco-2-ACE2 cells were infected and examined at multiple time points to observe infection kinetics. In all three cell lines, WT SARS-CoV-2-nLuc plateaued at 48 h post-infection. ΔE SARS-CoV-2-nLuc infection, while similar to WT virus infection at early time points, showed attenuated infectivity starting at 24 h post-infection. Collectively, these data demonstrate that deletion of E from SARS-CoV-2 results in attenuated viral infectivity including in human mucosal epithelial cells.

### ΔE SARS-CoV-2 virus infection produces expected sgRNAs and does not recombine with trans-complemented coE

Attenuation rather than loss of infectivity of ΔE SARS-CoV-2 could be explained by restoration of E gene expression, compensatory mutations, or a lack of expression of other viral genes due to the E gene deletion. To confirm that the ΔE SARS-CoV-2-nLuc genome had not been altered through passaging, RNA was isolated from passage 3 (P3) virus and sequenced. The E gene deletion made in the ΔE SARS-CoV-2-nLuc BAC was still present in P3 virus (**Fig 4A**), and no additional mutations were observed in the sequenced region, which spanned from the last 437 bp of S through the end of ORF6.

**Figure 4.**
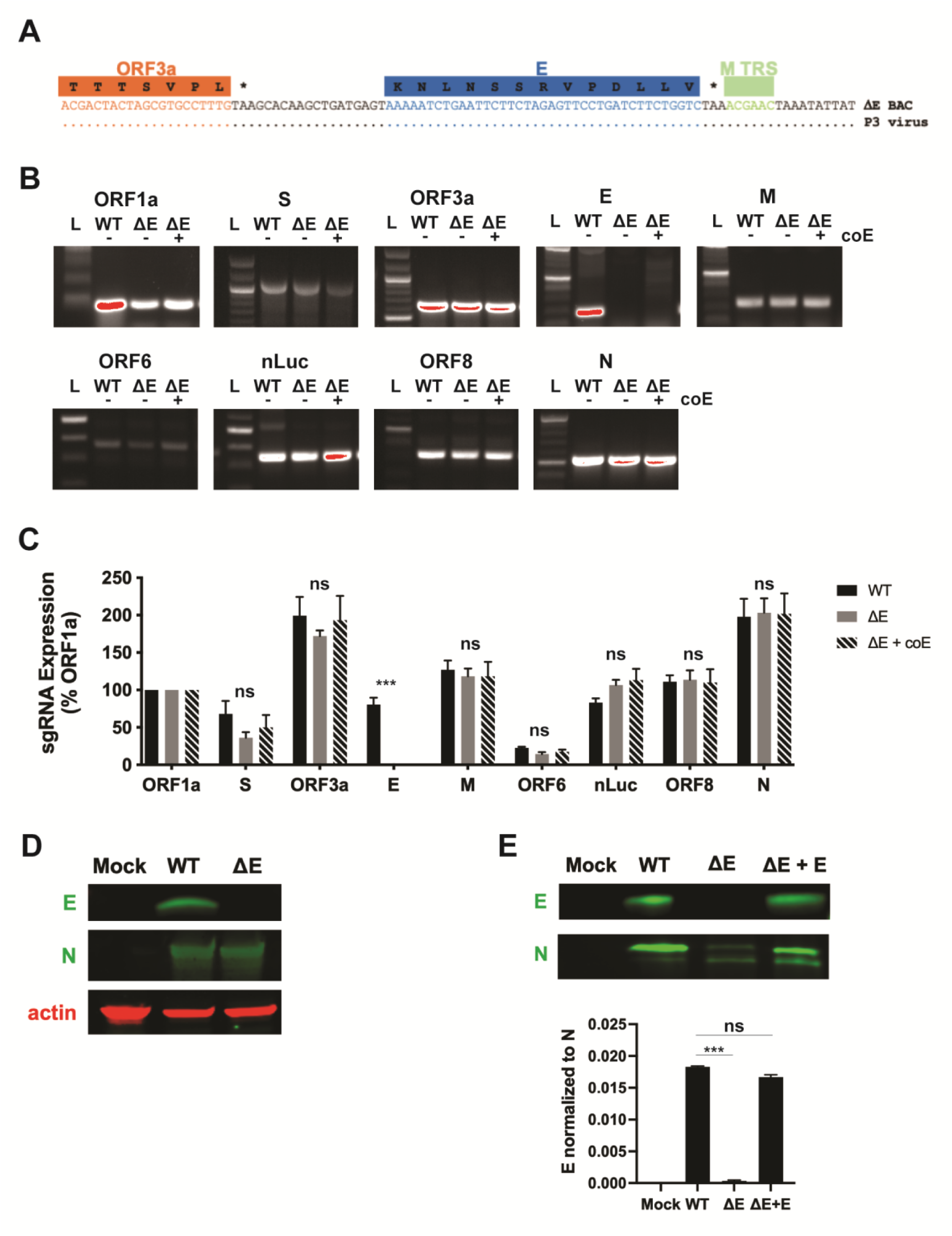
ΔE SARS-CoV-2 virus replication produces expected sgRNAs and packages E protein with trans-complemented coE. (A) The sequence of the E deletion of passage 3 virus (P3) is shown. (B) RNA was isolated from Vero E6-hAT (WT and ΔE) or E6-hAT cells stably expressing coE (ΔE only) infected with WT or ΔE SARS-CoV-2-nLuc virus. RT-PCR was performed with SARS-CoV-2 sgRNA specific primers and products were run on an agarose gel. L = 100 bp ladder. (C) Quantification of band intensity for sgRNAs was normalized to ORF1a. Statistical significance was determined by one-way ANOVA with Tukey’s multiple comparisons test. *** p < 0.001, ns p > 0.05. (D) Representative immunoblots of E, N, and actin expression in Vero E6-hAT cells infected with WT or ΔE SARS-CoV-2-nLuc are shown (n = 2). (E) Representative immunoblots of N and E in purified WT SARS-CoV-2-nLuc virions and ΔE SARS-CoV-2-nLuc virions produced with and without E expression are shown. Quantification of E normalized to N expression is graphed. *** p < 0.001, ns p > 0.05. Error bars represent SEM for 2 independent experiments.

To confirm that all sgRNAs except E are produced by ΔE SARS-CoV-2-nLuc, RNA from infected cells was isolated, RT-PCR amplified with sgRNA-specific primers, and visualized by agarose gel electrophoresis. For each virus, the amount of cDNA added to each RT-PCR reaction was normalized to ORF1a levels for each virus. All sgRNAs except E were detected in ΔE SARS-CoV-2-nLuc infected cells with similar relative abundance as for WT SARS-CoV-2-nLuc, with or without coE trans-complementation (**Fig 4B-C**). As expected, E sgRNA was not detected after ΔE SARS-CoV-2-nLuc infection with or without coE. In addition, RNA isolated from producer cells and supernatants from all three passages of WT SARS-CoV-2-nLuc or ΔE SARS-CoV-2-nLuc with and without coE was examined by qRT-PCR. While WT SARS-CoV-2-nLuc infection produced E sgRNA in cells and virus, no E sgRNA was detected with ΔE SARS-CoV-2-nLuc infection with or without coE (**Fig S4A**).

The lack of appreciable rescue of ΔE SARS-CoV-2-nLuc infectivity by coE trans-complementation could be due to a lack of coE expression, despite mRuby3 expression in induced coE cells. Enumeration of coE copies by qRT-PCR demonstrated that coE was expressed across all three passages in trans-complemented cells, albeit at a level 2- to 4-log lower than endogenous E detected during WT SARS-CoV-2-nLuc infection (**Fig S4B**, left). Expression of coE protein in induced Vero E6-hAT-coE cells was confirmed by immunoblot (**Fig S3C**), and the level of coE mRNA expression was similar in Vero E6-hAT-coE cells with or without infection by WT or ΔE SARS-CoV-2-nLuc (data not shown). Importantly, coE was not detected in ΔE SARS-CoV-2-nLuc virus produced in coE-expressing cells (**Fig S4B**, right), further confirming a lack of recombination of coE mRNA with the ΔE SARS-CoV-2-nLuc viral genome.

As sgRNA expression may not correlate with protein expression, E expression was measured in virions. As expected, while WT SARS-CoV-2-nLuc infection of Vero E6-hAT cells led to E protein expression, ΔE SARS-CoV-2-nLuc infection did not express E (**Fig 4D**). This resulted in E packaging in WT virions but not ΔE virions (**Fig 4E**). In contrast, ΔE SARS-CoV-2 produced in Vero E6-hAT-coE cells packaged E protein in virions similar to WT virions produced in Vero E6-hAT cells (**Fig 4E**), demonstrating that E trans-complementation was successful at the protein level.

### ΔE SARS-CoV-2 is defective for virus production and spread and is partially rescued by E trans-complementation

While ΔE SARS-CoV-2-nLuc infectivity was not significantly rescued by coE trans-complementation (**Fig 2A-C**), other characteristics of this mutant virus were rescued by E expression. While performing plaque titration of the viruses, we observed that ΔE SARS-CoV-2-nLuc generated small, pinprick-like plaques compared to large, distinct WT plaques (**Fig 5A-B**). In contrast, titration in cells expressing E resulted in ΔE SARS-CoV-2-nLuc plaques that matched WT plaques regardless of whether they were produced in cells expressing E. Similar results were observed with SARS-CoV-2 virus lacking the nLuc reporter gene (**Fig S2B**). Of note, it did not matter if ΔE SARS-CoV-2-nLuc was originally produced in cells expressing E, as all ΔE SARS-CoV-2-nLuc viruses produced small plaques when inoculated onto cells lacking E. Similarly, infecting Vero E6-hAT cells (lacking E expression) with ΔE SARS-CoV-2-nLuc produced in cells with or without E expression showed no appreciable difference in infectivity (**Fig S5A**). Conversely, ΔE SARS-CoV-2-nLuc infection of Vero E6-hAT cells expressing E resulted in 20-fold increased infectivity compared with ΔE SARS-CoV-2-nLuc infection of Vero E6-hAT cells without E (**Fig S5B**).

**Figure 5.**
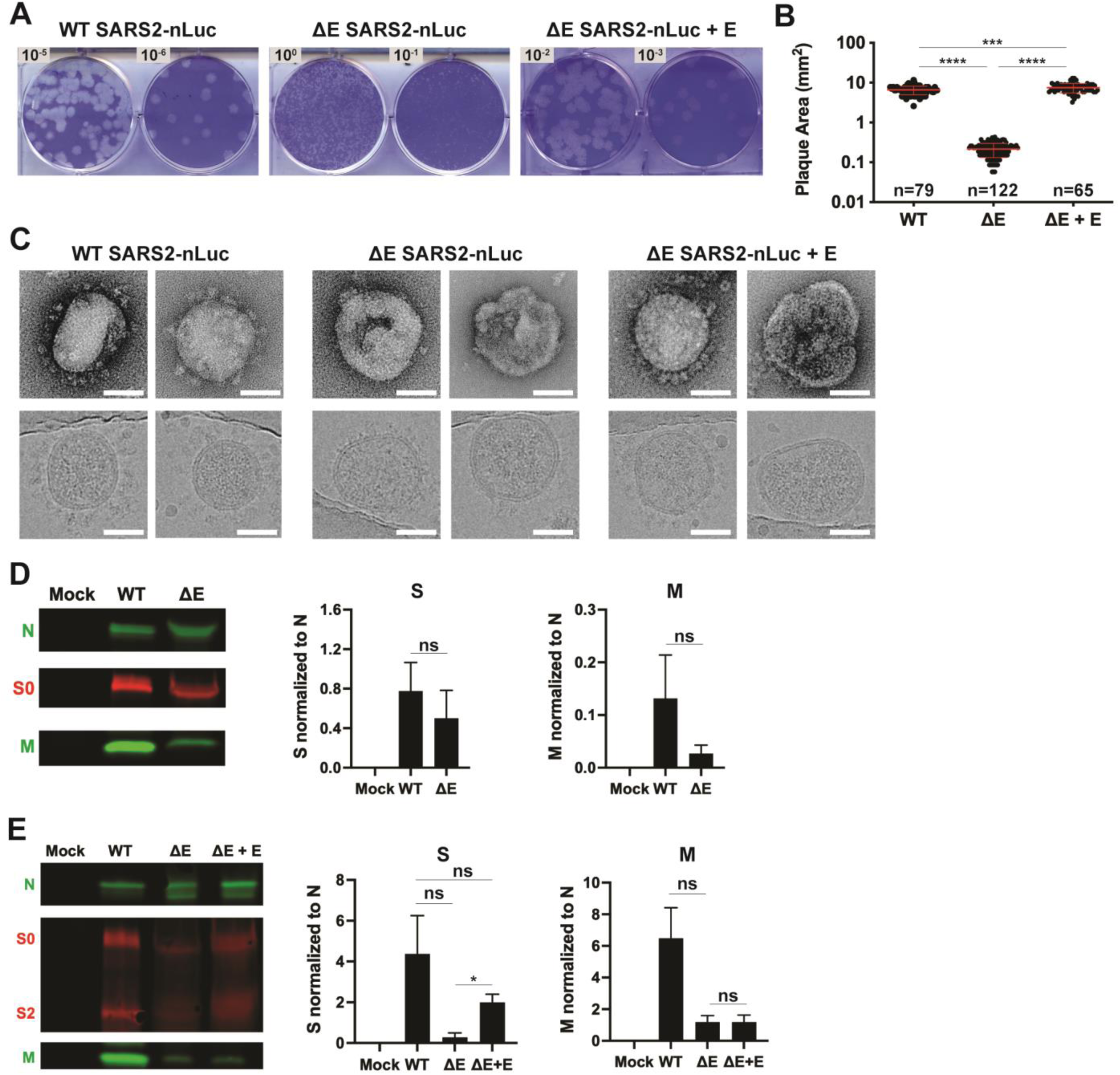
ΔE SARS-CoV-2 is defective for virus particle production and is partially rescued by E trans-complementation. (A) Plaque assays are shown for WT SARS-CoV-2-nLuc infection of Vero E6-hAT cells and ΔE SARS-CoV-2-nLuc infection of Vero E6-hAT cells with or without stable expression of E. Numbers represent log dilution of virus. (B) Plaque titer plates for viruses shown in (A) were digitized with a flatbed scanner and individual plaques were identified and measured with ImageJ to determine plaque area. Number of plaques examined are indicated for each virus. Statistical significance was determined by one-way ANOVA with Tukey’s multiple comparisons test. **** p < 0.0001, *** p < 0.001. Line represents mean and error bars represent standard deviation. (C) Negative stain EM (top) and Cryo-EM (bottom) of WT and ΔE SARS-CoV-2-nLuc produced with and without trans-complementation of E. Representative micrographs of two independent experiments are shown. Solid white bars represent 50 nm. (D) Representative immunoblots of N, S, and M expression in Vero E6-hAT cells infected with WT or ΔE SARS-CoV-2-nLuc are shown. Quantifications of S and M normalized to N expression are graphed. (E) Representative immunoblots of N, S, and M in purified WT SARS-CoV-2-nLuc virions and ΔE SARS-CoV-2-nLuc virions produced with and without E expression are shown. Quantification of S and M normalized to N expression are graphed. * p < 0.05, ns p > 0.05. Error bars represent SEM for three independent experiments.

In addition to having reduced virus particle production (**Fig 2B**), the ΔE SARS-CoV-2-nLuc particles differ from WT SARS-CoV-2-nLuc virus particles. While most WT virions visualized by negative stain EM and Cryo-EM were broadly decorated with S trimers, most ΔE SARS-CoV-2-nLuc virions displayed little or no S and often appeared misshapen (**Fig 5C**). Production of ΔE SARS-CoV-2-nLuc in cells expressing E resulted in a mixed phenotype, with some WT-like particles but also a population of defective particles similar to those produced in cells without E expression. While a similar amount of S was produced in cells infected with WT or ΔE SARS-CoV-2-nLuc (**Fig 5D**), purified viruses showed less incorporation of S in ΔE SARS-CoV-2-nLuc virions compared to WT virions, which could be partially rescued when E was supplied *in trans* and incorporated into virions at a level similar to WT virus (**Fig 5E**). Notably, S expression in infected cells and in WT virions was quite variable. Interestingly, less M was produced in cells infected with ΔE SARS-CoV-2-nLuc than WT virus, leading to less M incorporation in ΔE SARS-CoV-2-nLuc virions compared to WT (**Fig 5D-E**). This is consistent with a previous report showing that SARS-CoV-2 E RNA degradation by overexpression of the host protein RNF5 subsequently led to decreased M protein levels in infected cells (51) .

Examining nLuc expression of cells infected with WT or ΔE SARS-CoV-2-nLuc virus by immunofluorescence microscopy revealed that WT and ΔE viruses propagate differently. Vero E6-hAT, 16HBE-ACE2, and Caco-2-ACE2 cells infected with WT SARS-CoV-2-nLuc showed some contiguous patches of nLuc-expressing cells that were suggestive of cell-to-cell spread, as has been previously reported (52, 53), but predominantly consisted of scattered individual cells or small numbers of cells expressing nLuc that were consistent with particle-based spread (**Fig 6**). In contrast, cells from all three cell lines infected with ΔE SARS-CoV-2-nLuc showed little evidence of particle-based spread but instead contained large islands of nLuc-expressing cells that appeared to be syncytia. Interestingly, nuclear staining of these islands of nLuc-expressing cells showed a distinct “eye of the hurricane” pattern of aggregated nuclei with an empty center, which was not observed with WT SARS-CoV-2-nLuc infection. ΔE SARS-CoV-2-nLuc infection of E-expressing cells produced an increase in particle-based spread but also appreciably enhanced cell-to-cell transmission (**Fig 6A**). Similarly, WT SARS-CoV-2-nLuc infection of cells expressing E also showed an increase in cell-to-cell spread. Comparable results were observed with WT and ΔE SARS-CoV-2 lacking the nLuc reporter gene (**Fig S2E**). Collectively these data demonstrate that ΔE SARS-CoV-2 is defective for virus particle production and particle-based spread, is transmitted primarily cell-to-cell, and is partially rescued by E trans-complementation.

**Figure 6.**
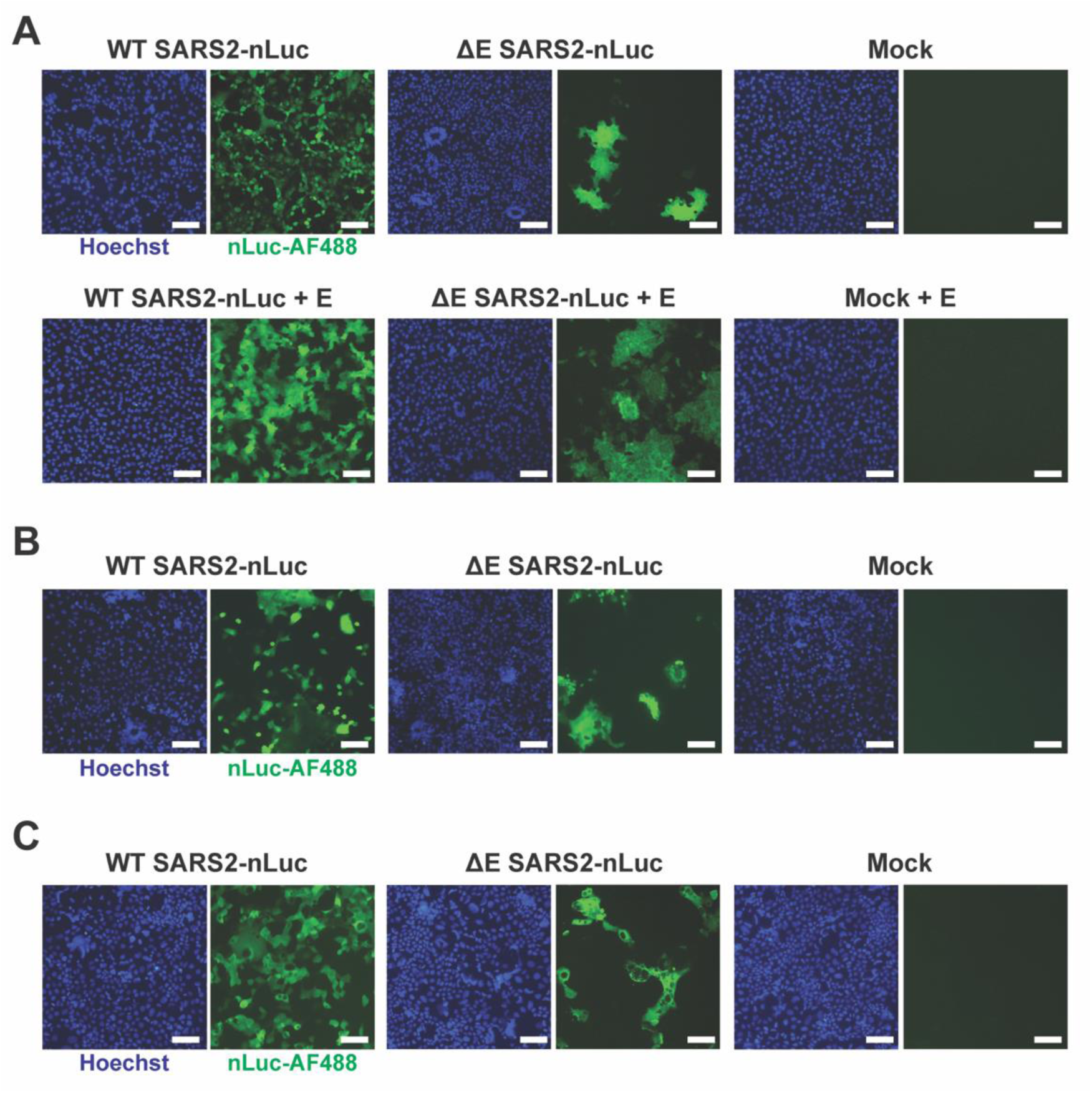
ΔE SARS-CoV-2 is defective for virus particle spread and is partially rescued by E trans-complementation. (A) Vero E6-hAT cells with or without stable expression of E were infected with WT or ΔE SARS-CoV-2-nLuc for 24 h, stained for nLuc, and imaged by widefield epifluorescence microscopy. Representative micrographs of three independent experiments are shown. Solid white bars represent 100 μm. (B) 16HBE-ACE2 and (C) Caco-2-ACE2 cells were infected with WT or ΔE SARS-CoV-2-nLuc for 24 h, stained for nLuc, and imaged by widefield epifluorescence microscopy. Representative micrographs of two independent experiments are shown. Solid white bars represent 100 μm.

### S neutralizing antibody inhibits SARS-CoV-2 virus particle spread but not cell-to-cell transmission

Neutralizing antibodies (NAbs) directed against S can effectively inhibit SARS-CoV-2 infection (54, 55). Protection of viral infections by NAbs appears to be predominantly based on inhibiting cell-free virus spread and targeting surface-exposed antigen on infected cells (55), which is circumvented by cell-to-cell virus transmission (53, 56–58) Since ΔE SARS-CoV-2 appeared to spread primarily cell-to-cell, we examined whether a NAb could prevent ΔE SARS-CoV-2 infection. The LY-CoV555 (bamlanivimab) NAb was effective at neutralizing USA-WA1/2020 SARS-CoV-2 (59–61) with an effective concentration to reduce infection by 50% (EC_50_) of 0.01 μg/mL in Vero E6-hAT cells (61). Vero E6-hAT cells were infected with WT or ΔE SARS-CoV-2-nLuc that had been pre-incubated for 1 h with NAb at the reported EC_50_, EC_90_, or 10 times the EC_90_ dose or left untreated. After 24 h the cells were assessed for nLuc activity (**Fig 7A**). The lowest dose of NAb was sufficient to reduce WT SARS-CoV-2-nLuc infectivity 10-fold, with no nLuc activity detected at the higher doses. In comparison, ΔE SARS-CoV-2-nLuc infectivity was reduced only 2-fold at the lowest dose and nLuc activity was still detectable at the higher doses, albeit reduced 100- and 500-fold (**Fig 7A**).

**Figure 7.**
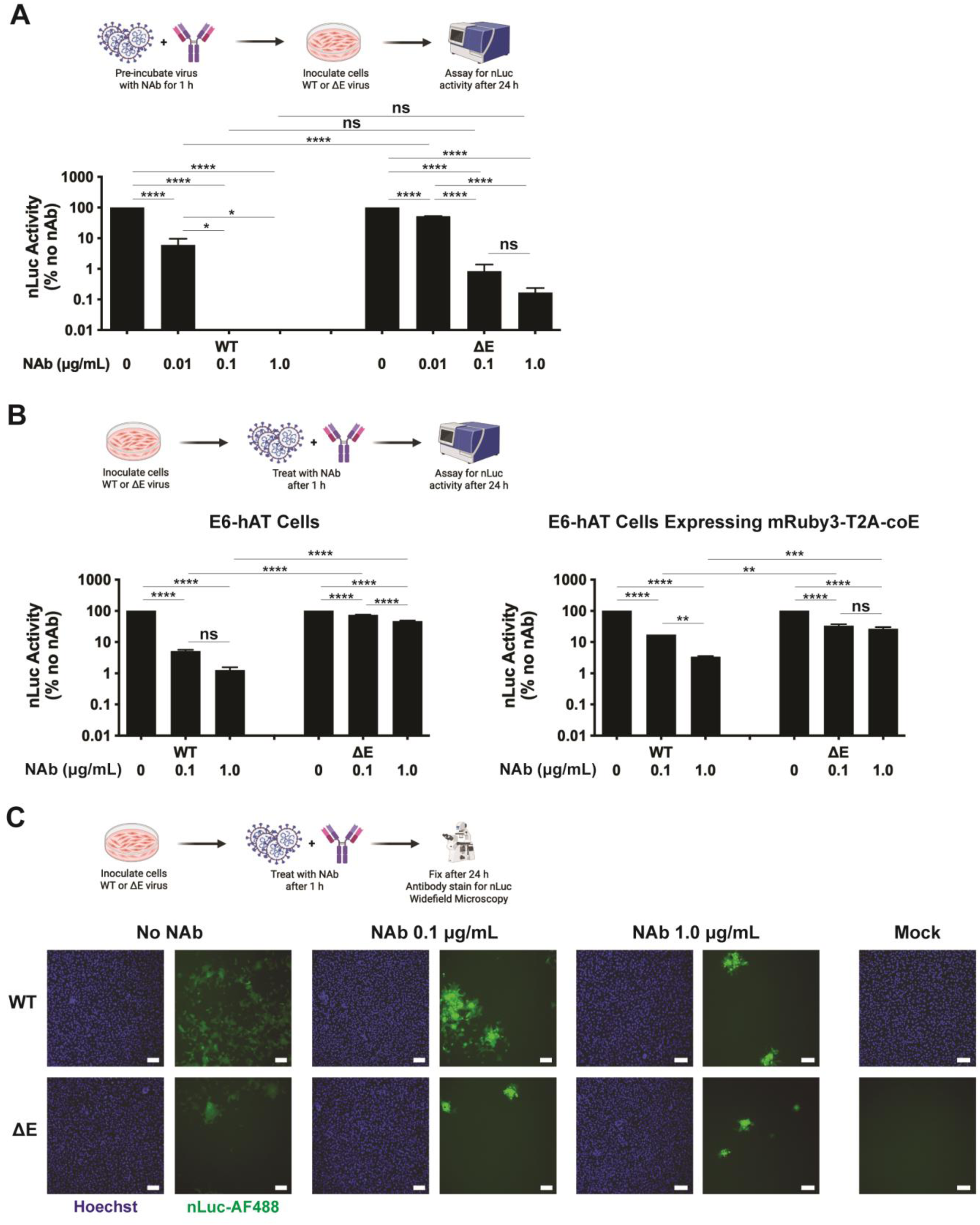
S NAb inhibits SARS-CoV-2 virus particle spread but not cell-to-cell transmission. (A) Vero E6-hAT cells were inoculated with WT or ΔE SARS-CoV-2-nLuc virus, which was pretreated with NAb for 1h, and assayed for nLuc activity after 24 h. Results for each virus are normalized to the no NAb condition. Error bars represent SEM for two independent experiments. (B,C) Vero E6-hAT cells were inoculated with WT or ΔE SARS-CoV-2-nLuc virus for 1h, treated with NAb, and after 24 h (B) assayed for nLuc activity or (C) antibody stained for nLuc and imaged by widefield epifluorescence microscopy. Results for each virus are normalized to the no NAb condition. Statistical significance was determined by two-way ANOVA with Tukey’s multiple comparisons test. **** p < 0.0001, *** p < 0.001, ** p < 0.01, * p < 0.05, ns p > 0.05. Error bars represent SEM for 2-3 independent experiments. Representative micrographs of two independent experiments are shown. Solid white bars represent 100 μm.

To directly compare WT and ΔE SARS-CoV-2 cell-to-cell transmission, a method was needed to permit initial infection of cells but to inhibit subsequent cell-free infection while allowing cell-to-cell spread. Thus, Vero E6-hAT cells were infected with WT or ΔE SARS-CoV-2-nLuc for 1 h before treatment with NAb and assayed for nLuc activity after 24 h (**Fig 7B**). As the EC_50_ dose of NAb did not inhibit WT or ΔE SARS-CoV-2-nLuc when applied post-inoculation (data not shown), only the EC_90_ and 10 times EC_90_ doses were used. Like what was observed with NAb pre-treatment, WT SARS-CoV-2-nLuc infectivity was reduced 10- and 100-fold compared to no NAb treatment, whereas ΔE SARS-CoV-2-nLuc infectivity was only reduced 1.5- and 3-fold (**Fig 7B**, left). Similar results were seen in Vero E6-hAT cells expression coE (**Fig 7B**, right).

We hypothesized that the difference in NAb efficacy at inhibiting WT and ΔE SARS-CoV-2-nLuc infectivity after virus challenge was due to the relative contribution of virus particle spread to overall infectivity for each virus but that both viruses retained the ability to transmit cell-to-cell. To confirm this, nLuc expression was visualized in cells with and without NAb pre-treatment and infected with viruses (**Fig 7C**). Without NAb treatment, both WT and ΔE SARS-CoV-2-nLuc exhibited similar patterns of predominantly cell-to-cell or cell-free spread as shown in **Fig 6A**. Treating with increasing doses of NAb inhibited particle-based spread but preserved cell-to-cell spread by both WT and ΔE SARS-CoV-2-nLuc (**Fig 7C**). At the highest dose of NAb (1.0 μg/mL, 10 times EC_90_) no particle-based spread was apparent for either WT or ΔE SARS-CoV-2-nLuc. Altogether, these results demonstrate that NAb treatment inhibits SARS-CoV-2 virus particle spread but not cell-to-cell transmission and that ΔE SARS-CoV-2 is transmitted primarily cell-to-cell.

*SARS-CoV-2-mediated cell fusion is enhanced by E expression in infected target cells* Inhibiting SARS-CoV-2 virus particle spread while preserving cell-to-cell spread through high dose NAb treatment affords the ability to directly compare WT and ΔE SARS-CoV-2 cell-to-cell transmission without the confounding effects of potentially unequal virus particle spread. While the pattern of infected cells during high dose NAb treatment suggests transmission is occurring cell-to-cell via syncytia formation (**Fig 7C**), it is also possible that localized virus particle spread without cell fusion occurs. To determine whether SARS-CoV-2 transmission with high NAb concentrations occurs through syncytia formation, we employed a bimolecular fluorescence complementation (BiFC) cell fusion reporter system, in which the linked Renilla luciferase (rLuc) and enhanced green fluorescent protein (eGFP) genes are split into N- and C-terminal fragments, NrLuc-eGFP and rLuc-eGFPC (62). Stable expression of either the NrLuc-eGFP or rLuc-eGFPC fragment was accomplished by transducing Vero E6-hAT cells with a lentiviral vector encoding one of the fragments, generating Vero E6-hAT-N and Vero E6-hAT-C cell lines. Both lines were co-cultured 1:1, infected with SARS-CoV-2-nLuc at a low multiplicity of infection (MOI=0.01) for 1 h, and treated with NAb for 24 h. Then the cells were fixed and imaged for nLuc expression by fluorescence microscopy (**Fig S6A**). Fusion of adjacent Vero E6-hAT-N and Vero E6-hAT-C cells resulted in the reconstitution of rLuc and eGFP and produced green fluorescence. Infected cells, which were positive for nLuc expression, were also positive for reconstituted eGFP expression, indicating that SARS-CoV-2 transmission in the presence of a high NAb concentration occurs through cell fusion (**Fig S6B**).

To compare cell fusion induced by WT and ΔE SARS-CoV-2-nLuc, Vero E6-hAT cells with or without E expression were infected with WT or ΔE virus for 1 h, treated with NAb for 24 h, fixed, and stained for nLuc expression (**Fig 8A**). Infection with WT SARS-CoV-2-nLuc resulted in larger syncytia than ΔE SARS-CoV-2-nLuc in Vero E6-hAT cells (**Figs 8B, S6B**). Syncytia size was enhanced for both viruses in Vero E6-hAT-coE cells expressing E. Interestingly, the “eye of the hurricane” aggregated nuclei structures observed with ΔE SARS-CoV-2-nLuc infection without NAb treatment (**Fig 6A**) were also present during WT SARS-CoV-2-nLuc infection with NAb treatment (**Fig 8A**). Collectively, these data suggest virus-mediated cell fusion is enhanced by E expression in infected target cells.

**Figure 8.**
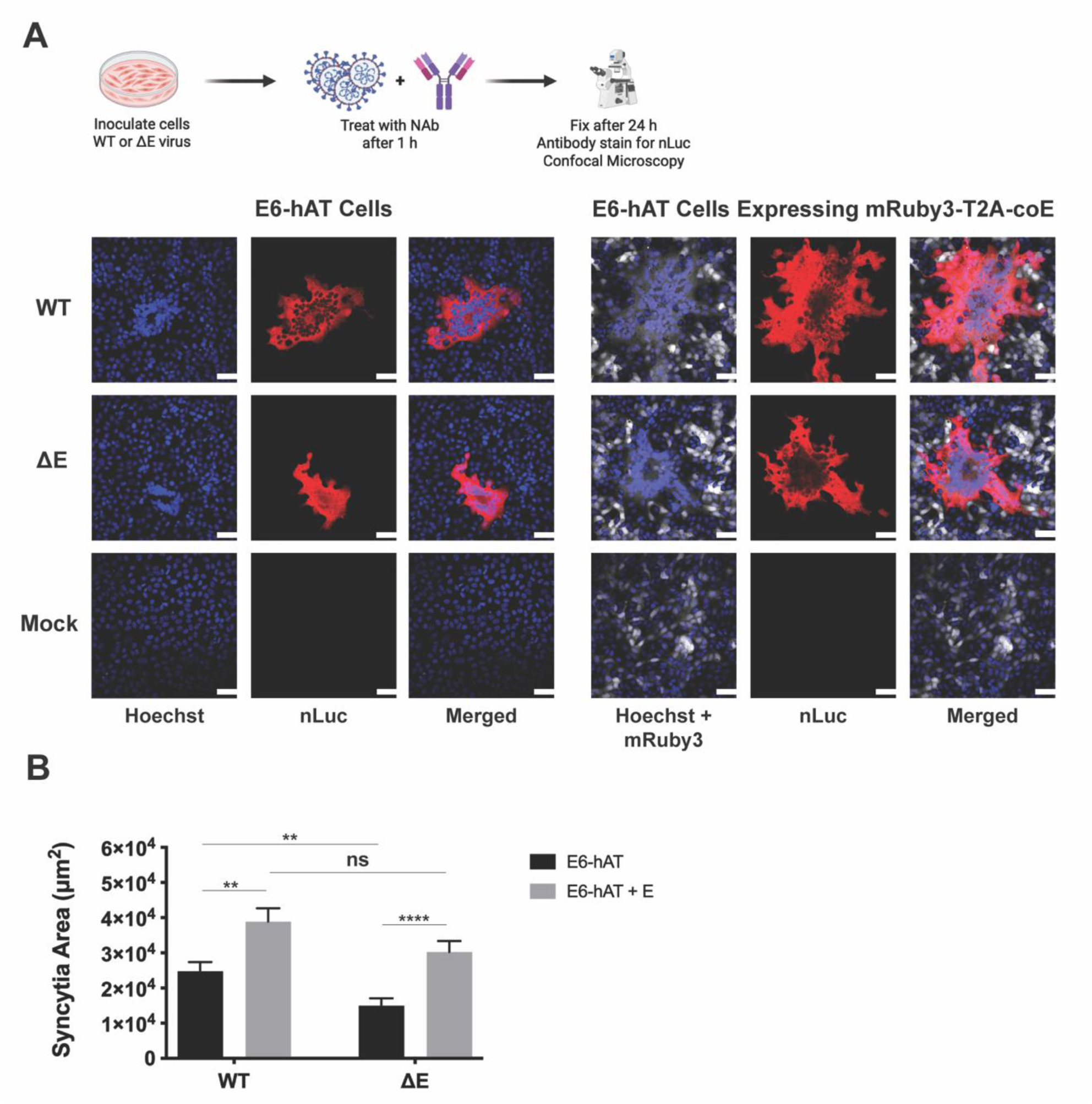
SARS-CoV-2-mediated cell fusion is enhanced by E expression in infected target cells. (A,B) Vero E6-hAT cells with and without stable E expression were infected with WT or ΔE SARS-CoV-2-nLuc virus (MOI=0.01) for 1 h, treated with Nab for 24 h, and stained with Hoechst (blue) and nLuc (red). mRuby3 expression was also assessed (white). Representative confocal micrographs (A) and quantification of 35-40 total fields per condition (B) are shown. Solid white bars represent 50 μm. **** p < 0.0001, *** p < 0.001, ** p < 0.01, * p < 0.05, ns p > 0.05. Error bars represent SEM for four independent experiments.

### Lack of E during SARS-CoV-2 infection results in greater S cell surface expression and ERGIC reorganization

Prior work has demonstrated that expression of SARS-CoV-2 S protein in cells results in its expression on the cell surface (37, 63). In contrast, co-expressing E with S causes S to be predominantly retained within the ERGIC to facilitate virus particle assembly (37). Given our observation that fewer S trimers were incorporated into ΔE SARS-CoV-2-nLuc virus particles compared with WT particles (**Fig 5C,E**), we hypothesized that the lack of E expression during ΔE SARS-CoV-2-nLuc infection results in less S retention within the ERGIC and increased expression at the cell surface. To examine S localization during WT and ΔE SARS-CoV-2-nLuc infection, Vero E6-hAT cells were infected with WT or ΔE SARS-CoV-2-nLuc and fixed after 24 h. The cells were stained prior to permeabilization with anti-S primary and red fluorescent secondary antibodies to detect cell surface expression. After permeabilization, the cells were stained with anti-S primary and green fluorescent secondary antibodies to detect total (surface and internal) expression. After imaging via confocal microscopy, green fluorescence that co-localized with red fluorescence (representing cell surface expressed S that was labeled with both sets of antibodies) was masked to visualize distinct populations of surface (red) and internal (green) S (**Fig 9A**). Infection with ΔE SARS-CoV-2-nLuc resulted in increased S surface expression compared with WT virus (**Fig 9B**). While the proportion of surface-expressed S in ΔE SARS-CoV-2-nLuc-infected cells was highly variable, there was nonetheless a trend for higher S surface expression in some cells infected with ΔE SARS-CoV-2-nLuc compared to WT virus.

**Figure 9.**
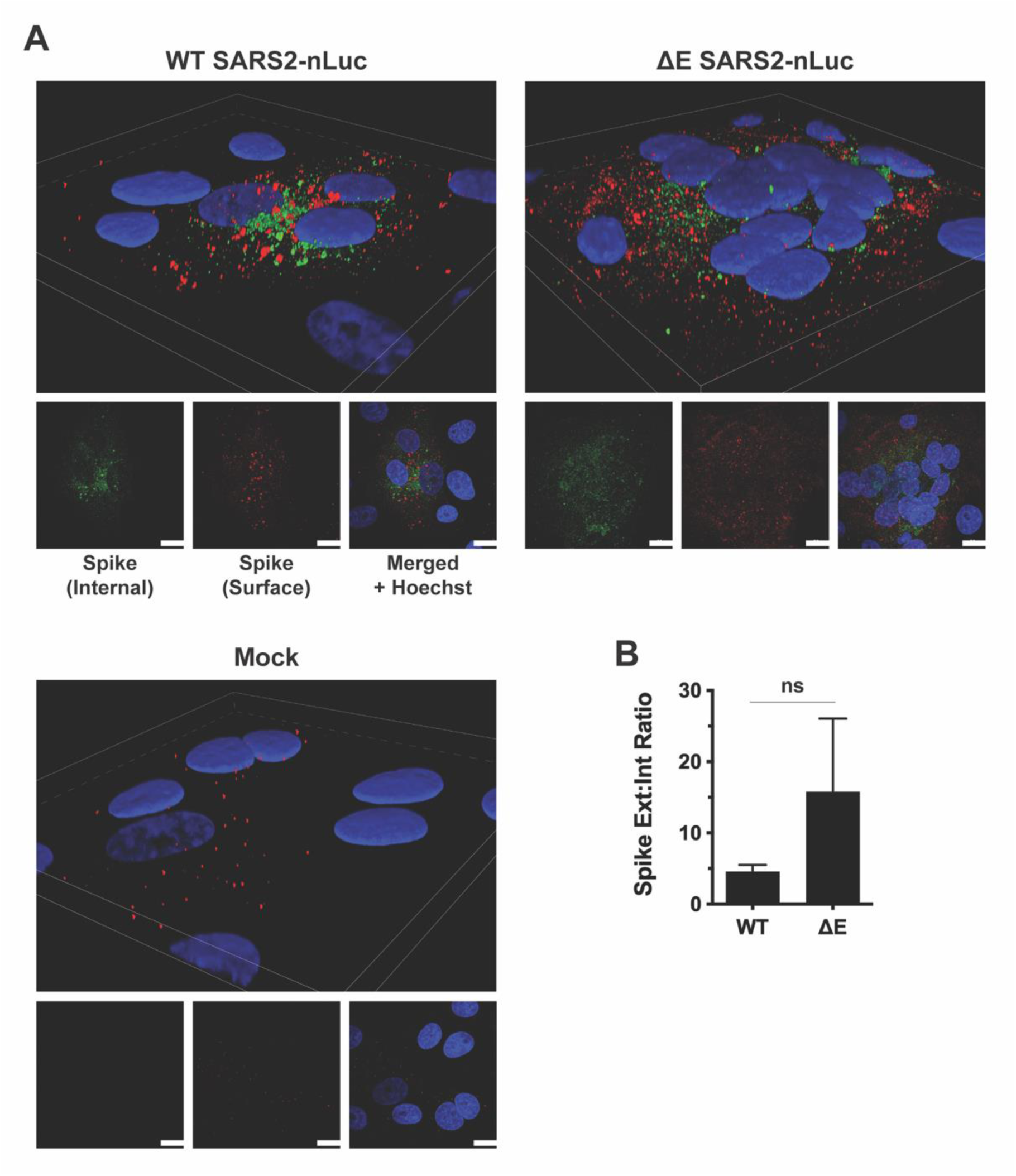
Lack of E during SARS-CoV-2 infection results in greater S cell surface expression. (A,B) Vero E6-hAT cells were infected with WT or ΔE SARS-CoV-2-nLuc virus for 24 h and stained for surface (red) and internal (green) S expression and Hoechst (blue). Representative confocal micrographs (A) in volume (top) and maximum intensity (bottom) projections are shown. Quantification of internal and external S was performed and graphed as a ratio (B). Error bars represent SEM for two independent experiments. Solid white bars represent 20 μm.

Different localization patterns of internal S in cells infected with WT or ΔE SARS-CoV-2-nLuc were observed (**Fig 9A**). While most internal S in cells infected with WT SARS-CoV-2-nLuc appeared in centralized, perinuclear structures consistent with ERGIC localization, S inside ΔE SARS-CoV-2-nLuc-infected cells was more diffuse, raising the question of whether S in cells infected with ΔE SARS-CoV-2-nLuc was retained within the ERGIC. To test this, Vero E6-hAT cells were infected with WT or ΔE SARS-CoV-2-nLuc for 24 h and stained for S and the canonical ERGIC marker protein ERGIC-53 (**Fig 10A**). Cells infected with WT virus exhibited the same ERGIC-53 staining pattern as uninfected cells, with S co-localizing with ERGIC-53. In contrast, cells infected with ΔE

**Figure 10.**
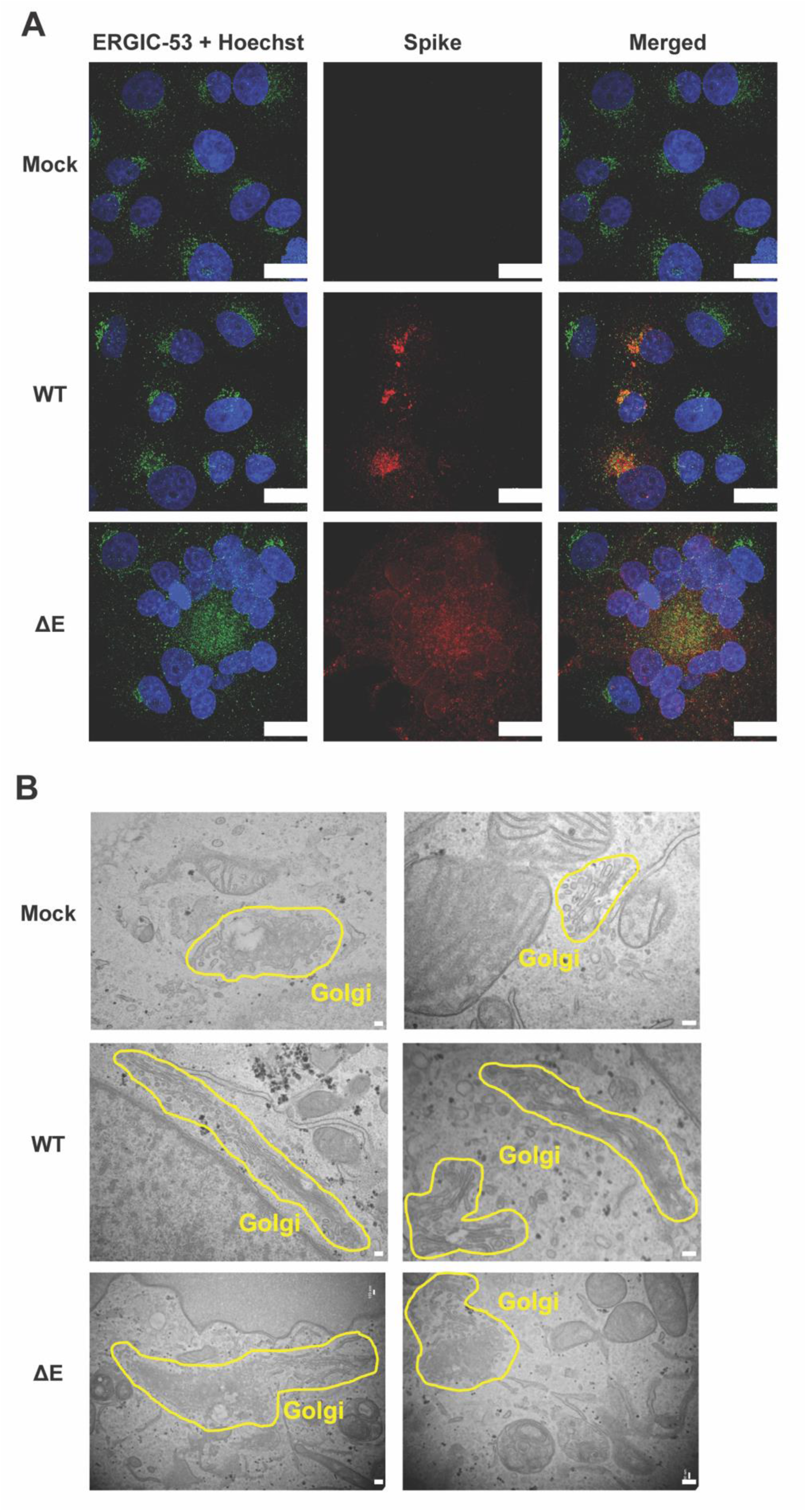
Lack of E during SARS-CoV-2 infection results in ERGIC reorganization. (A) Vero E6-hAT cells were infected with WT or ΔE SARS-CoV-2-nLuc for 24 h, stained for ERGIC-53 (green), S (red) and Hoechst (blue), and imaged by confocal microscopy. Solid white bars represent 20 μm. (B) Vero E6-hAT cells were infected with WT or ΔE SARS-CoV-2-nLuc for 24 h and imaged by transmission EM. Solid white bars represent 200 nm.

SARS-CoV-2-nLuc also exhibited strong co-localization of S and ERGIC-53, but the ERGIC structure was reorganized, appearing more diffuse compared to uninfected cells or cells infected with WT virus. Within ΔE virus-mediated syncytia, the center of the “eye of the hurricane” aggregated nuclei structure was filled with diffuse ERGIC-53 co-localized with S. Similar results were observed in Vero E6-hAT cells stably expressing E (**Fig S7**). Induced mRuby3 expression was frequently observed to have lower expression in infected cells, especially within syncytia. To directly observe structural changes in the ERGIC during ΔE SARS-CoV-2-nLuc infection, Vero E6-hAT cells infected with WT and ΔE SARS-CoV-2-nLuc were examined by transmission electron microscopy. Cells infected with WT virus exhibited similar Golgi structure as mock infected cells, although the Golgi appeared elongated (**Fig 10B**). In contrast, infection with ΔE SARS-CoV-2-nLuc resulted in fragmentation and reorganization of the Golgi similar to that observed with the ERGIC by immunofluorescence microscopy.

To determine if this altered ERGIC organization was due to SARS-CoV-2-induced syncytia or specific to ΔE SARS-CoV-2 infection, syncytia were directly compared in cells infected with WT or ΔE SARS-CoV-2-nLuc (**Fig 11A**). Unlike the diffuse ERGIC observed with ΔE SARS-CoV-2-nLuc syncytia, WT virus syncytia retained multiple dense ERGIC structures. Thus, ERGIC/Golgi reorganization is not induced by syncytium formation but rather by a lack of E expression by the virus.

**Figure 11.**
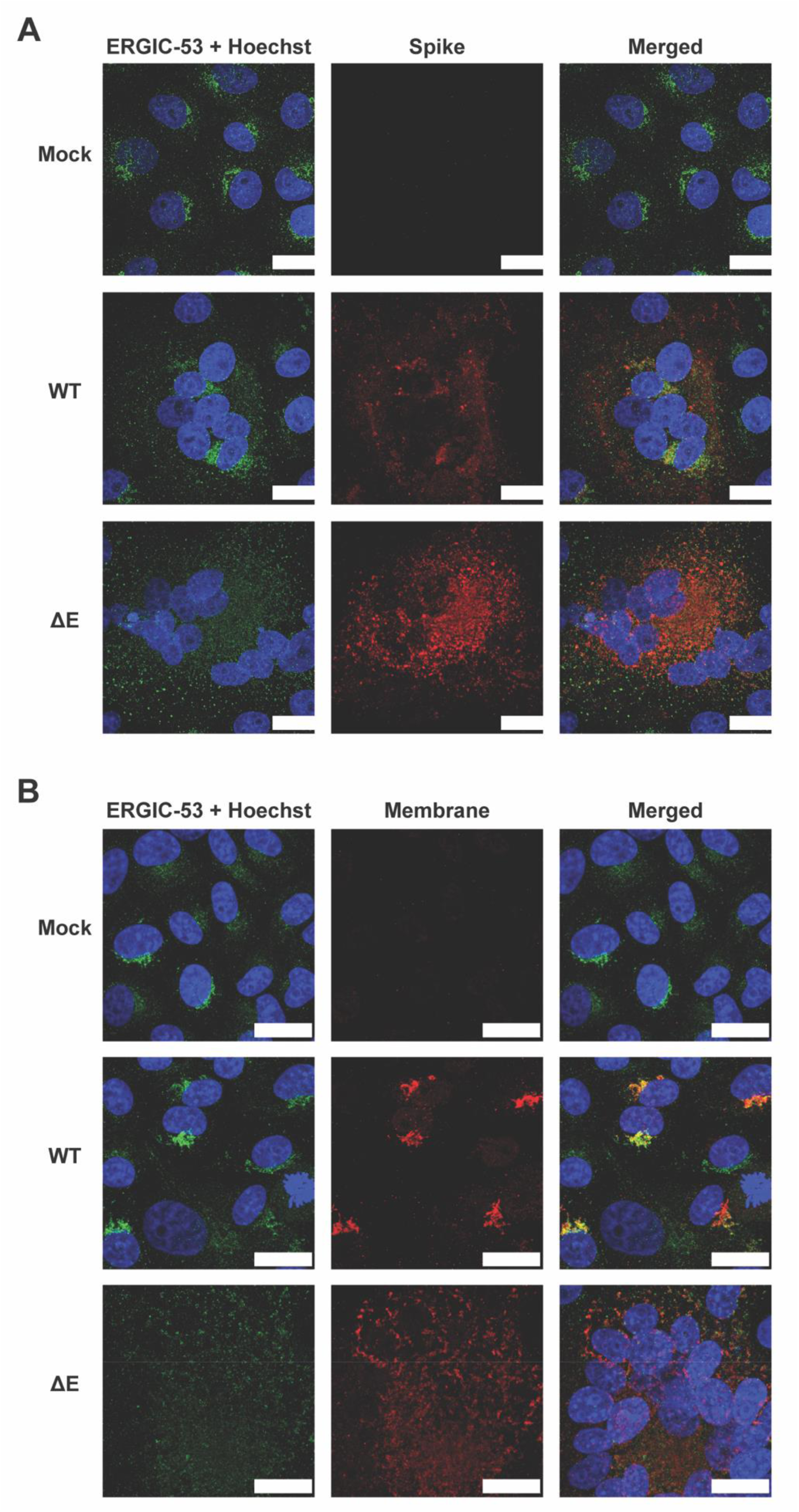
ERGIC is not reorganized in WT SARS-CoV-2 syncytia and M localizes to the ERGIC regardless of reorganization. (A) Vero E6-hAT cells were infected with WT or ΔE SARS-CoV-2-nLuc for 24 h, stained for ERGIC-53 (green), S (red) and Hoechst (blue), and imaged by confocal microscopy. (B) Vero E6-hAT cells were infected with WT or ΔE SARS-CoV-2-nLuc for 24 h, stained for ERGIC-53 (green), M (red) and Hoechst (blue), and imaged by confocal microscopy. Solid white bars represent 20 μm.

As M has been shown to also localize to the ERGIC and associate with E, M localization was examined in Vero E6-hAT cells infected with WT or ΔE SARS-CoV-2-nLuc. As observed with S, M localized with the ERGIC (**Fig 11B**). Again, WT SARS-CoV-2-nLuc infection did not dramatically alter ERGIC-53 staining, which was co-localized with M. Although ΔE SARS-CoV-2-nLuc induced ERGIC fragmentation, M was still colocalized with ERGIC-53.

To determine whether ERGIC reorganization was specific to Vero E6 cells, 16HBE-ACE2 cells were infected with WT or ΔE SARS-CoV-2-nLuc and stained for S and ERGIC-53. As with Vero E6-hAT cells, WT SARS-CoV-2-nLuc infection did not alter ERGIC staining compared to uninfected cells. However, ΔE SARS-CoV-2-nLuc infection resulted in ERGIC reorganization similar to what was observed in Vero E6-hAT cells (**Fig 12**). Altogether, these data demonstrate that a lack of E during SARS-CoV-2 infection results in greater S cell surface expression and reorganization of the ERGIC and Golgi.

**Figure 12.**
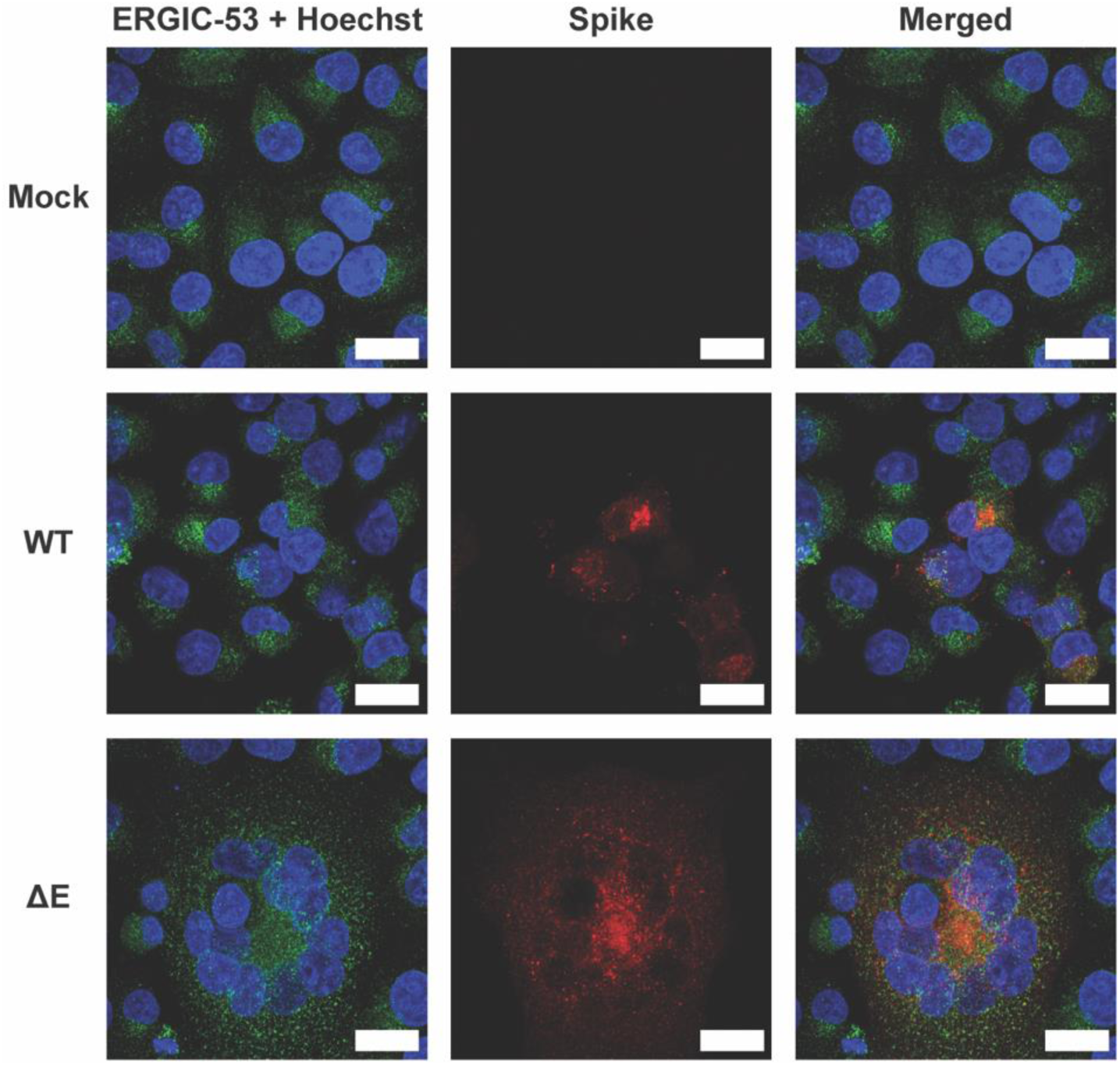
ERGIC reorganization during ΔE SARS-CoV-2 infection occurs in human respiratory epithelial cells. 16HBE-ACE2 cells were infected with WT or ΔE SARS-CoV-2-nLuc for 24 h, stained for ERGIC-53 (green), S (red) and Hoechst (blue), and imaged by confocal microscopy. Solid white bars represent 20 μm.

## Discussion

Previous studies have shown that TGEV, HCoV-OC43, and MERS-CoV lacking E are unable to replicate (40–42). In contrast, MHV and SARS-CoV can replicate without E but are attenuated (43, 44), which we also observed for ΔE SARS-CoV-2-nLuc in Vero-E6-hAT cells and multiple human epithelial cell lines. We showed that ΔE SARS-CoV-2-nLuc attenuation was due to reduced infectious particle production and increased cell-to-cell transmission (**Fig 13**). Plaque titration of ΔE SARS-CoV-2-nLuc demonstrated fewer and much smaller plaques compared to the large, distinct plaques observed with WT virus, which was also observed for MHV and SARS-CoV (43, 44). Increased cell surface expression of S was observed for ΔE SARS-CoV-2-nLuc, which led to significantly reduced virus production and release from the ERGIC pathway and any virions that were produced had little incorporation of S. This was expected given the previously reported roles of CoV E in virion assembly and release (35–37). In addition, transfection studies showed that SARS-CoV-2 E expression slows down the secretory pathway, inducing retention of glycoproteins in the ERGIC, including S (37). We hypothesized that a lack of intracellular S expression could lead to its expression at the cell surface, which was true to an extent, but surface S localization was variable. Unexpectedly, ΔE SARS-CoV-2-nLuc infection led to changes in ERGIC and Golgi morphology.

**Figure 13.**
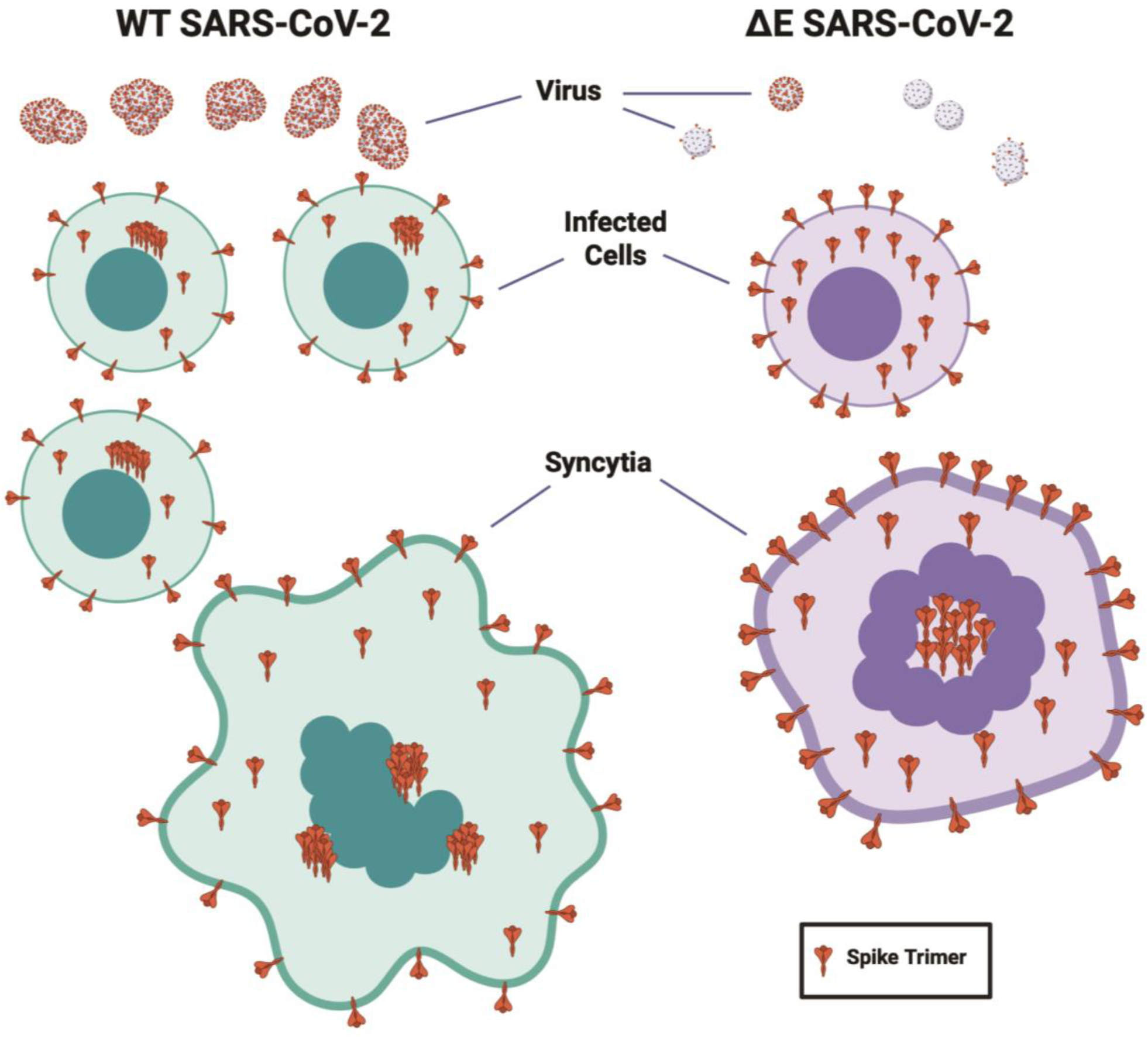
Model schematic comparing WT and ΔE SARS-CoV-2 replication. Differences between WT and ΔE SARS-CoV-2 virus particle production and S incorporation, cell surface expression and intracellular localization of S, and syncytia structure are shown.

WT SARS-CoV-2-nLuc exhibited reduced infectivity in 16HBE cells compared with ΔE SARS-CoV-2-nLuc, which was opposite of the relative infectivity observed between WT and ΔE SARS-CoV-2-nLuc in all other cell lines. As overexpression of ACE2 in 16HBE cells rescued the WT SARS-CoV-2-nLuc infectivity defect, this would suggest ΔE SARS-CoV-2 infectivity is less sensitive to ACE2 expression than WT virus. This could be explained by our observation that while WT virus spreads via cell-to-cell or cell-free transmission, ΔE SARS-CoV-2 spreads primarily cell-to-cell. While cell-to-cell spread of SARS-CoV-2 is dependent upon S expression (53, 64), the expression of ACE2 is not strictly necessary (65, 66). Expression levels of other host cell factors, such as TMPRSS2, could also impact relative WT and ΔE SARS-CoV-2 infectivity. Other studies have measured TMPRSS2 expression in these cell lines and found the highest expression level in Calu-3 cells, with Caco-2 and Vero E6 cells having 40% and 6% the level of TMPRSS2 in Calu-3 cells, respectively (67), and 16HBE cells having 20-fold less TMPRSS2 than Calu-3 cells (68). The combination of low ACE2 and TMPRSS2 expression in 16HBE cells coupled with ACE2-independent cell-to-cell virus spread could permit ΔE SARS-CoV-2 to infect 16HBE cells better than WT virus. 16HBE cells could be a useful tool in further exploring how ΔE SARS-CoV-2 infection differs from WT SARS-CoV-2.

ΔE SARS-CoV-2-nLuc infectivity and virus particle production and transmission by trans-complementation of E was only partially rescued, despite similar levels of E incorporation into virus particles as observed with WT SARS-CoV-2-nLuc. One factor in this limited rescue is the expression level of trans-complemented E, which had 2-3-log lower mRNA abundance in infected cells versus endogenous E sgRNA during WT virus infection. While SARS-CoV-2 has been shown to inhibit host protein synthesis during translation and mRNA nuclear export (69–71), the abundance of trans-complemented E mRNA was similar in uninfected and infected cells, suggesting that the virus did not inhibit mRNA production or stability. However, cells trans-complemented with E had lower mRuby3 protein expression after SARS-CoV-2-nLuc infection, particularly within syncytia, suggesting virus-induced host protein inhibition reduced trans-complemented E protein expression levels. Another factor limiting the efficacy of E trans-complementation could be timing of expression. While E is expressed during SARS-CoV-2 infection, in trans-complemented cells E is present prior to infection and at lower levels based on RNA abundance. This different timing and level of expression could alter how E influences virus replication. Additionally, E undergoes post-translational modification, including palmitoylation, during SARS-CoV-2 infection (35, 72). Trans-complemented E could have altered post-translational modifications and/or trafficking, impacting one or more if its functions. Optimization of the trans-complementation system to overcome these limitations may improve rescue of ΔE SARS-CoV-2.

The variability in E dependency across CoVs suggests E performs multiple roles in virus assembly and release (73), some of which are crucial and others that may be dispensable or that can be substituted for by other viral proteins (74). We confirmed that attenuation of ΔE SARS-CoV-2 was authentic and not due to reconstitution of the E gene, emergence of compensatory mutations, or the lack of ORF7a/b in our nLuc reporter viruses. Examining production of sgRNAs by ΔE virus with and without E trans-complementation showed that all sgRNAs except E sgRNA are produced with similar abundance to WT. This demonstrates that attenuated ΔE SARS-CoV-2 infectivity is not due to a defect in viral RNA synthesis but suggests the replication defect occurs after RNA synthesis. While deleting either the E gene or the ORF3 gene in SARS-CoV reduced but did not eliminate viral titers, deletion of both E and ORF3a abrogated SARS-CoV and SARS-CoV-2 replication, indicating that ORF3a can substitute for some functionality of E (45, 46). This would make both viral proteins potential candidates for compensatory mutations to overcome replication defects introduced by the deletion of E. However, no mutations were observed after three passages of ΔE SARS-CoV-2.

While higher surface expression of S during ΔE SARS-CoV-2-nLuc infection was anticipated to promote increased cell fusion compared to WT infection, we observed greater cell fusion and larger viral plaque size with WT SARS-CoV-2-nLuc infection than with ΔE virus infection. We also observed that trans-complementation of E enhanced cell fusion and viral plaque size, including with WT SARS-CoV-2-nLuc infection, suggesting that E is also needed for optimum cell fusion. The C-terminus of E contains a PSD-95/Dlg/ZO-1 (PDZ)-binding motif (PBM), facilitating interactions of E with various PDZ-containing host proteins, including Zona Occludens-1 (ZO-1) and Proteins Associated with Lin Seven 1 (PALS1), which are key regulators of tight junction (TJ) formation and epithelial monolayer integrity (75, 76). Further, E expression has been shown to downregulate expression of TJ proteins Occludin and Claudin-4 (77), and deletion or mutation of the E PBM has been shown to produce smaller viral plaques, reduced disruption of ZO-1 in tight junctions, and overall reduced pathogenicity (78). Disruption of TJs and epithelial monolayer integrity by E could facilitate cell fusion. Alternatively, the viroporin activity of E, by altering Ca^2+^ homeostasis, could promote syncytia formation, which has been shown to be dependent upon Ca^2+^ oscillations (64, 79, 80). Thus, while increased S surface expression with ΔE SARS-CoV-2 infection could promote greater cell fusion, the lack of E expression results in overall reduced cell fusion, possibly due to the absence of the viroporin activity of E or its regulation of tight junctions. Further studies are needed to elucidate how E expression enhances virus-induced cell fusion.

It is generally agreed that CoV assembly occurs in or near the ERGIC but trafficking of nascent virions from the ERGIC to extracellular space is less certain. Different pathways may be utilized and/or altered by different CoVs, including the Golgi secretory pathway, the endosome recycling compartment, or lysosomal exocytosis (81–84). Fragmentation of the Golgi and its translocation to the center of virus-induced syncytia was previously observed with MHV (85). Fluorescence imaging also noted Golgi fragmentation during late SARS-CoV-2-nLuc infection, when M, S, and E proteins were expressed and after recruitment of the lysosomal marker LAMP1 to the Golgi/ERGIC, likely due to fusion of lysosomes with ERGIC membranes (86) . However, we only observed ERGIC fragmentation after infection with ΔE SARS-CoV-2-nLuc, suggesting that this phenomenon is transient during late SARS-CoV-2 infection and is enhanced if E is not present.

SARS-CoV-2 infection has been shown to result in an accumulation of autophagosomes but inhibition of cargo degradation through reduced autophagic flux, processes which involve E and M and which may lead to virus particle release (87, 88). The localization of M to ERGIC-derived autophagosomes is dependent upon expression of E, without which M localizes to the Golgi (89–91). It could be envisioned that without E, accumulation of M – and perhaps nascent virions – in the Golgi/ERGIC could promote the observed fragmentation and translocation, as well as alter the pathway through which virions are released. As E has been described in slowing down the secretory pathway (37), its absence could accelerate virus-induced processes within the secretory compartments. Additional studies investigating SARS-CoV-2 virus particle assembly and trafficking are needed to examine how the presence or absence of E alters these processes as well as the cellular secretory system.

## Materials and Methods

### Biosafety statement

All SARS-CoV-2 virus production and infections were performed at biosafety level 3 in the Regional Biocontainment Laboratory at the University of Pittsburgh Center for Vaccine Research.

### SARS-CoV-2 BAC

The SARS-CoV-2 USA-WA1/2020 viral genome was obtained as 7 individual cDNA fragments cloned within pUC57 or pCC1BAC via a Materials Transfer Agreement from the University of Texas Medical Branch (48), with and without replacement of the ORF7a/b gene by the mNeonGreen (mNeon) reporter gene. The mNeon reporter gene in fragment 7 was replaced via Gibson cloning with the codon-optimized nLuc gene from a reporter lentiviral genome (pNL4-nLucCO-6ATRi-BAL) that we had previously produced (92).

SARS-CoV-2 genome fragments 1-4 and fragments 5-7 were each assembled using the Golden Gate Assembly kit per the manufacturer’s instructions (NEB), with type IIB restriction enzymes BsaI-HFv2 and Esp3I, respectively. A MEC was produced via gene synthesis (GenScript), consisting of the CMV and T7 promoters upstream of the first 351 bp of the SARS-CoV-2 USA-WA1/2020 genome (from the start of the 5’ UTR up to the unique BsiWI restriction site within the nsp1 gene) and the last 1,398 bp of the viral genome (from the unique XhoI restriction site within the nucleocapsid gene through the end of the 3’ UTR) followed by a polyA(29) tail, HDV ribozyme, and bovine growth hormone polyadenylation signal sequence. This MEC was cloned into pCC1BAC via NotI/BamHI restriction cloning to produce a mammalian expression BAC (pBAC), which was subsequently digested with BsiWI and XhoI and incubated with digested fragments 1-4 (BsiWI restriction enzyme) and fragments 5-7 (XhoI restriction enzyme) together with T4 DNA ligase to produce the full SARS-CoV-2 genome with and without an nLuc reporter. To facilitate assembly of the full-length SARS-CoV-2 genome into the BAC, the 5’ end (the first 351 bp of the genome consisting of the 5’-UTR and the N-terminal of the nsp1 gene through the unique BsiWI restriction site) and the 3’ end (the last 1,398 bp of the genome from the unique XhoI restriction site in nucleocapsid through the 3’-UTR) of the viral genome were synthesized as part of the MEC. The full SARS-CoV-2 genome sequence within the BAC was verified by Sanger sequencing.

### ΔE SARS-CoV-2 BAC

The E TRS and the first 186 bp of the E gene were deleted from the SARS-CoV-2 genome to produce ΔE SARS-CoV-2 that lacked expression of E but preserved the expression of the downstream M gene and all subsequent genes. Briefly, portions of fragments 6 and 7 from UTMB were PCR amplified and Topo cloned into pCR2.1-TOPO (ThermoFisher Scientific). Deletion of E was performed by Gibson cloning. The fragments were then assembled as described above. The full ΔE SARS-CoV-2 genome sequence within the BAC was verified by Sanger sequencing.

### Inducible SARS-CoV-2 E expression constructs

pLVX-TetOne-Puro-mRuby3co-T2A-coE was produced by gene synthesis (GenScript) of the codon-optimized versions of mRuby3 and the SARS-CoV-2 USA-WA1/2020 E gene linked by the T2A peptide, which was cloned by EcoRI/AgeI restriction cloning into the pLVX-TetOne-Puro-hAXL plasmid (Addgene). A variation replacing the puromycin resistance gene in the pLVX-TetOne-Puro vector with the NeoR/KanR/G418 resistance gene from pIRES (Takara Bio) via Gibson cloning produced pLVX-TetOne-G418-mRuby3co-T2A-coE.

### Cell fusion/syncytia reporter constructs

The N- and C-terminal fragments of the split renilla luciferase and enhanced green fluorescent protein fusion (Split-rLuc-eGFP) BiFC reporter were cloned from pCG-NrLuc-eGFP and pCG-rLuc-eGFPC, respectively (62), into the XbaI/SalI digested pLenti-CMV-Hygro-ACE2 plasmid (Addgene) via Gibson cloning to produce pLenti-CMV-Hygro-NrLuc-eGFP and pLenti-CMV-Hygro-rLuc-eGFPC.

### Cells

Vero E6 cells (ATCC) are AGM (*Cercopithecus aethiops*) kidney cells, BHK cells (ATCC) are Syrian hamster (*Mesocricetus auratus*) kidney cells, HEK 293T and 293F cells (ATCC) are human epithelial cells, Calu-3 cells (kind gift from Douglas Reed, University of Pittsburgh) are human lung adenocarcinoma cells, Caco-2 cells (ATCC) are human colorectal adenocarcinoma cells, and 16HBE14o- (16HBE) cells (kind gift from John Alcorn, University of Pittsburgh) are human bronchial epithelial cells. These cells were grown in Dulbecco’s Modified Eagle Medium (DMEM) containing 10% or 20% (Caco-2) fetal bovine serum (FBS; Atlanta Biologicals), 100 U/ml penicillin, 100 μg/ml streptomycin, and 2 mM L-glutamine (PSG; ThermoFisher Scientific), termed DMEM-10 medium, at 37° C and 5% CO_2_. 293F cells were maintained at 37° C with 5% CO_2_ in FreeStyle 293 Expression Medium (Thermo Fisher) supplemented with penicillin and streptomycin (Gibco). Vero E6-TMPRSS2-T2A-ACE2 cells (BEI Resources) referred to herein as Vero E6-hAT cells, were grown in DMEM-10 supplemented with 10 μg/ml puromycin (Invivogen).

16HBE-ACE2 and Caco-2-ACE2 cells were produced by stable transduction of 16HBE and Caco-2 cells, respectively, with pLenti-CMV-Hygro-ACE2 lentivirus and maintained in DMEM-10 or DMEM-20 supplemented with 500 μg/ml hygromycin (Invitrogen). Vero E6-hAT-NrLuc-eGFP (Vero E6-hAT-N) and Vero E6-hAT-rLuc-eGFPC (Vero E6-hAT-C) cells were produced by stable transduction of Vero E6-hAT cells with pLenti-CMV-Hygro lentivirus encoding the N-terminal or C-terminal fragment of the Split-rLuc-eGFP fusion reporter gene, respectively, and maintained in DMEM-10 supplemented with 10 μg/ml puromycin and 500 μg/ml hygromycin.

BHK-coE cells were produced by stable transduction of BHK cells with pLVX-TetOne-Puro lentivirus encoding mRuby3-T2A-coE and maintained in DMEM-10 supplemented with 3 μg/ml puromycin. Vero E6-hAT-coE cells were produced by stable transduction of Vero E6-hAT cells with pLVX-TetOne-G418-mRuby3-T2A-coE lentivirus and maintained in DMEM-10 supplemented with 10 μg/ml puromycin and 500 μg/ml G418 (Gibco). Vero E6-hAT-N-coE and Vero E6-hAT-C-coE cells were produced by stable transduction of Vero E6-hAT-N and Vero E6-hAT-C cells, respectively, with pLVX-TetOne-G418-mRuby3-T2A-coE lentivirus and maintained in DMEM-10 supplemented with 10 μg/mL puromycin, 500 μg/ml hygromycin, and 500 μg/ml G418. TetOne-mediated expression of mRuby3-T2A-coE was induced with 100 ng/ml doxycycline (Thermo Scientific) for 48 h prior to use, which was maintained throughout the course of the experiment. Expression of mRuby3-T2A-coE was visually confirmed by widefield epifluorescence microscopy prior to use.

### Production of lentiviruses

pLenti and pLVX viruses were produced by Lipofectamine 2000 (ThermoFisher Scientific) transfection in 293T cells with the pCAGGS-PAX2 lentiviral packaging plasmid and the pL-VSV-G plasmid. Virus was harvested 48 h later, centrifuged at 2000 x g for 20 m, filtered through a 0.45 μm polyethersulfone (PES) filter (Millipore), aliquoted, and frozen at -80° C.

### Production of SARS-CoV-2 virions

Transfection of 20 μg viral cDNA into 4 x 10^6^ BHK cells in a 100 mm dish was performed with Lipofectamine 2000 or Polyethylenimine (PEI, Sigma-Aldrich). The medium was removed after 24 h and the cells were split 1:2, overlaid onto two 100 mm dishes each of 5 x 10^6^ Vero E6-hAT cells, and incubated for 3 days. Supernatants were centrifuged at 2000 x g for 20 m, filtered through a 0.45 μm PES or mixed cellulose ester (MCE) filter, and aliquoted frozen at -80° C or lysed in TRIzol-LS (ThermoFisher Scientific). For ΔE SARS-CoV-2 virus production, transfections were also performed in BHK-coE and Vero E6-hAT-coE cells induced with doxycycline.

### nLuc assay

nLuc from transfected or infected cells was measured by lysing cells in Nano-Glo Luciferase Assay Buffer (Promega), incubation with Nano-Glo Luciferase Assay Substrate (Promega), and detection on a BioTek Synergy2 Multi-Detection Microplate Reader.

### RT-PCR

RNA was isolated from TRIzol-LS-treated virus or TRIzol-treated cell lysates as previously described (93). RT-PCRs were performed using primers shown in **Table 1** and PCR reactions were run on an agarose gel for visualization. Band intensities were quantified using a Bio-Rad Gel Doc EZ Imager and Image Lab software. Quantitative RT-PCRs were performed using primers and probes listed in **Table 1** with plasmids used as standards and run on a Bio-Rad CFX96 Real-Time PCR System.

### Plaque assay

Vero E6-hAT cells were plated at 1 x 10^6^ cells/well overnight in 6-well plates. Virus was diluted in 10-fold serial dilutions in OptiMEM medium (ThermoFisher Scientific) + 2% FBS. Medium was removed from cells and 200 μl virus dilutions were added in duplicate to wells and incubated at 37° C, 5% CO_2_ for 1 h, with plates tilted every 15 m. The virus was removed from all wells and cells were washed once with phosphate buffered saline (PBS, Gibco) before overlay with 3 ml of 1.25% Avicel (Millipore Sigma) in EMEM medium (Quality Biological) with 10% FBS, PSG, and Non-Essential Amino Acids (Gibco). Plates were incubated 2-5 days and the Avicel was removed in PBS. Cells were fixed in 10% formalin for 24 h at room temperature (RT). Wells were washed with PBS and stained with 0.2% crystal violet in 25% methanol for 15 m at RT. Stained plates were washed in water to remove excess stain and allowed to air dry overnight. Plaques were imaged on a flatbed scanner and counted.

### Virus particle EM

SARS-CoV-2 virions were fixed in equal volume 4% paraformaldehyde (PFA) with 10 mM HEPES for 48 h at 4° C, ultracentrifuged through a 20% sucrose cushion in NTE buffer (100 mM NaCl, 10 mM Tris-HCl, 1 mM EDTA, pH 7.6) for 2.5 h, 4° C at 117,250 x g, and resuspended in NTE buffer. For negative stain EM, 3 μL of sample was applied to a freshly glow-discharged continuous carbon copper grid and stained with 1% uranyl acetate solution. The grids were then inserted into a Tecnai TF20 electron microscope (Thermo Fisher Scientific, MA, USA), equipped with a field emission gun and images captured using an XF416 CMOS camera (TVIPS GmbH, Gilching, Germany) to assess particle assembly and presence of Spike. For Cryo-EM, 3 μL of sample was applied to a freshly glow-discharged Quantifoil R2/1 grid (Quantifoil Micro Tools GmbH, Großlöbicha, Germany) and then blotted and plunge-frozen with a Vitrobot Mk 4 (TFS). Grids were mounted on a Krios 3Gi cryo-electron microscope operating at 300 kV and equipped with a Falcon 4i direct electron detecting camera (TFS). Images were collected in electron counting mode under the control of the TFS EPU software using a total dose of ∼30 e/Å^2^. A magnification of 85,000x was used, correspond to 1.51 Å per pixel at the sample.

### Western blot

Unless otherwise indicated below for specific experiments, standard immunoblot sample preparation was as follows: cells were lysed in 1X Laemmli sample buffer (Bio-Rad) containing 2-mercaptoethanol (Sigma-Aldrich) and transferred to fresh microcentrifuge tubes. Lysates were denatured at 100 °C for 3 m and briefly centrifuged. Samples were loaded onto pre-cast 4-15% TGX gradient polyacrylamide gels (Bio-Rad) and subjected to SDS-PAGE gel electrophoresis at 70 V for 2 h. Proteins were transferred to nitrocellulose membranes (Millipore) using a wet electroblotting system (Bio-Rad) in 1X transfer buffer prepared from 5X stock (124 mM Tris, 960 mM glycine, 0.05% SDS) supplemented with 20% methanol, and run at 75 V for 1 h at 4 °C. Membranes were washed in Tris-buffered saline (TBS, Boston BioProducts), blocked in Intercept TBS blocking buffer (LI-COR) for 1 h at RT, and incubated with primary antibodies diluted in Intercept TBS blocking buffer supplemented with 0.2% Tween-20 (T-20 diluent) overnight at 4 °C. Membranes were washed three times for 5 m each with TBS supplemented with 0.2% Tween-20 (TBS-T), incubated with secondary antibodies (donkey α-mouse IRDye 680LT and donkey α-rabbit IRDye 800CW, LI-COR, 1:10,000) diluted in T-20 diluent for 1 h at RT, and washed three times for 5 m with TBS-T before a final rinse with TBS. Membranes were imaged on a LI-COR Odyssey Clx imager using ImageStudio software with default acquisition settings, and band intensities were quantified using the ImageStudio application.

For coE expression, BHK-coE and Vero E6-hAT-coE cells were seeded at 2 x 10^6^ cells/dish in 100 mm dishes with and without 250 ng/mL doxycycline. After 48 h, cells were washed once with PBS and prepared according to the above standard protocol with primary antibodies against E (rabbit α-E, GTX636915, GeneTex, 1:1,000) and actin (goat α-actin, sc-1616, Santa Cruz Biotechnology, 1:1,000) and secondary antibodies (donkey α-goat IRDye 680LT and donkey α-rabbit IRDye 800CW, LI-COR, 1:10,000).

For virus-infected cells, Vero E6-hAT cells were seeded at 5 x 10^5^ cells/well in 6-well plates and infected with WT virus (MOI=0.1) or ΔE virus (MOI=0.3), or mock infected. After 12 h, cells were washed once with PBS and prepared according to the above standard protocol with primary antibodies against S (mouse α-S, GTX632604, GeneTex, 1:1,000), M (rabbit α-M, PAB31758, Abnova, 1:500), E (rabbit α-E, GTX636915, GeneTex, 1:1,000), and N (rabbit α-N, 200-401-A50, Rockland, 1:3,000).

For virus supernatants, Vero E6-hAT and induced Vero E6-hAT-coE cells were seeded overnight at 15 x 10^6^ cells/dish in 150 mm dishes and infected with virus (MOI=0.01) or mock infected. After 48 h (WT virus) or 72 h (ΔE virus), supernatants were centrifuged at 2000 x g for 20 m, filtered through a 0.45 μm PES filter, supplemented with 10 mM HEPES, ultracentrifuged through a 20% sucrose cushion in NTE buffer for 2.5 h, 4 °C at 117,250 x g, resuspended overnight at 4 °C in 50 μl RIPA buffer (Boston BioProducts) supplemented with cOmplete protease inhibitor (Roche), alilquoted, and frozen at -20 °C. Concentrated supernatants were combined with 2X Laemmli sample buffer supplemented with 355 mM 2-Mercaptoethanol and denatured at 100 °C for 3 m. Samples were prepared according to the above standard protocol with primary antibodies against S (mouse α-S, GTX632604, GeneTex, 1:1,000), M (rabbit α-M, PAB31758, Abnova, 1:500), E (rabbit α-E, GTX636915, GeneTex, 1:1,000), and N (rabbit α-N, 200-401-A50, Rockland, 1:3,000).

### Neutralizing antibody assays

DNA encoding SARS-CoV-2 S NAb LY-CoV555 heavy- and light-chain variable domain genes were synthesized by IDT and cloned into a modified pVRC to produce full length human IgG1 antibodies as previously described (94) . Recombinant LY-COV555 was produced by polyethylenimine (PEI) facilitated, transient transfection of 293F cells that were maintained in FreeStyle 293 Expression Medium. Transfection complexes were prepared in OptiMEM and added to cells. Five days post-transfection supernatants were harvested, clarified by low-speed centrifugation, and incubated overnight with Protein Agarose Resin (GoldBio). The resin was collected in a chromatography column, washed with a column volume of 10 mM tris(hydroxymethyl)aminomethane-HCl (TRIS), 150 mM NaCl at pH 7.5, and eluted in 0.1M Glycine (pH 2.5) which was immediately neutralized by 1M TRIS, (pH 8.5). Antibodies were then dialyzed against PBS (pH 7.4). Vero E6-hAT cells were plated at 1.25 x 10^5^ cells/well overnight in 24-well plates. In one set of experiments, virus was pre-treated with NAb for 1 h at 37° C prior to infecting cells in duplicate wells for 24 h. In a second set of experiments, cells were infected with virus for 1 h before virus was removed from all wells and cells were washed once with PBS, fed with DMEM-10 supplemented with NAb, and incubated for 24 h. For both sets of experiments infection was measured by nLuc assay.

### Immunofluorescence microscopy

Unless otherwise indicated below for specific experiments, standard immunofluorescence microscopy sample preparation was as follows: Cells were seeded into 35 mm glass-bottom dishes (MatTek) and incubated until near confluence, overnight to 48 h. Cells were fixed with 4% PFA for 24 h at 4° C, permeabilized with 0.1% Triton X-100 (ThermoFisher Scientific) for 15 m at RT, blocked with 5% normal goat serum (Invitrogen) for 1 h at RT, incubated with primary antibody for 1 h at RT, incubated with secondary antibody for 1 h at RT, counterstained with Hoechst 33342 (1:2000, ThermoFisher Scientific) for 15 m at RT, and mounted with 1.5 glass coverslips. Blocking and immunofluorescence staining/wash buffers contained 0.5% bovine serum albumin (ThermoFisher Scientific) and 1% glycine (Millipore Sigma) in PBS.

For ACE2 cell surface expression, cells were counterstained with Hoechst 33342 prior to fixation with 2% PFA for 20 m at RT. No permeabilization was performed. Primary antibody (mouse α-ACE2 MAB9332, R&D Systems, 1:100) was incubated for 24 h at 4° C. Secondary antibody (goat α-mouse-AlexaFluor488, Invitrogen, 1:500) was incubated for 1 h at RT. Confocal laser scanning microscopy was performed on a Nikon A1 microscope equipped with a motorized piezo Z stage and 60X objective, and images were analyzed with Nikon Elements.

For nLuc expression, cells were seeded at 3 x 10^5^ cells/well overnight into either 6-well plates or MatTek dishes, infected with virus for 1 h, washed with PBS, and fed with DMEM-10, which in some experiments was supplemented with NAb. After 24 h, cells were fixed and prepared according to the above standard protocol with primary (mouse α-nLuc MAB10026, R&D Systems, 1:500) and secondary (goat α-mouse-AlexaFluor488, 1:500) antibodies. Widefield epifluorescence microscopy was performed on a Motic inverted fluorescence microscope with a 10X objective and images were analyzed with Fiji/ImageJ (95) .

For ERGIC-53 and S expression, cells were seeded at 1.5 x 10^5^ cells/dish overnight into MatTek dishes, infected with virus (MOI=0.01) for 1 h, washed with PBS, and fed with DMEM-10. After 24 h, cells were fixed and prepared according to the above standard protocol with primary (mouse α-ERGIC-53 C-6, Santa Cruz, 1:100 and rabbit α-S 40150-R007 Sino Biological, 1:500) and secondary (goat α-mouse-AlexFluor488, 1:500 and goat α-rabbit-AlexaFluor568, Invitrogen, 1:500) antibodies. Confocal laser scanning microscopy was performed on a Nikon A1 microscope equipped with a 60X objective and motorized piezo Z stage, and images were analyzed with Nikon Elements.

For ERGIC-53 and M expression, cells were seeded at 1.5 x 10^5^ cells/dish overnight into MatTek dishes, infected with virus (MOI=0.01) for 1 h, washed with PBS, and fed with DMEM-10. After 24 h, cells were fixed and prepared according to the above standard protocol with primary (mouse α-ERGIC-53 C-6, Santa Cruz, 1:100 and rabbit α-M PAB31758 Abnova, 1:250) and secondary (goat α-mouse-AlexFluor488, 1:500 and goat α-rabbit-AlexaFluor568, Invitrogen, 1:500) antibodies. Confocal laser scanning microscopy was performed on a Nikon A1 microscope equipped with a 60X objective and motorized piezo Z stage, and images were analyzed with Nikon Elements.

### Cell fusion assay

Vero E6-hAT and induced Vero E6-hAT-coE cells were seeded at 1 x 10^6^ cells/dish overnight in MatTek dishes, infected with virus (MOI=0.01) for 1 h, washed with PBS, and fed with DMEM-10 supplemented with NAb (1 μg/ml). After 24 h, cells were fixed and prepared according to the above standard protocol with primary (mouse α-nLuc MAB10026, 1:500) and secondary (goat α-mouse-AlexaFluor647, Invitrogen, 1:500) antibodies. Confocal laser scanning microscopy was performed on a Nikon A1 microscope with a 40X objective, with single focal plane images taken of 9-10 fields on average for four independent experiments. Images were analyzed and total area of each syncytium (as indicated by nLuc expression) was determined with Nikon Elements software.

### Split-eGFP BiFC syncytia assay

Vero E6-hAT-N and Vero E6-hAT-C cells with and without induced expression of coE were co-cultured 1:1 at 1 x 10^6^ total cells/dish overnight in MatTek dishes, infected with virus (MOI=0.01) for 1 h, washed with PBS, and fed with DMEM-10 supplemented with NAb (1 μg/ml). After 24 h, cells were fixed and prepared according to the above standard protocol with primary (mouse α-nLuc MAB10026, 1:500) and secondary (goat α-mouse-AlexFluor647, 1:500) antibodies. Confocal laser scanning microscopy was performed on a Nikon A1 microscope with a 40X objective, with single focal plane images taken of 6 fields each for two independent experiments. Images were analyzed with Nikon Elements software.

### S localization assay

Vero E6-hAT cells were seeded at 1.5 x 10^5^ cells/dish overnight in MatTek dishes, infected with virus (MOI=0.01) for 1 h, washed with PBS, and fed with DMEM-10. After 24 h, cells were fixed with 4% PFA for 24 h at 4° C, blocked with normal goat serum, incubated with primary (rabbit α-S 40150-R007, 1:500), and secondary (goat α-rabbit-AlexaFluor568, 1:500) surface antibodies, permeabilized, re-blocked, incubated with primary (rabbit α-S 40150-R007, 1:500) and secondary (goat α-rabbit-AlexaFluor488, Invitrogen 1:500) surface/intracellular antibodies, and counterstained with Hoechst 33342. Confocal laser scanning microscopy was performed on a Nikon A1 microscope equipped with a 60X objective and motorized piezo Z stage, with 4-6 Z-stacks taken for each of two independent experiments. Images were analyzed with Nikon Elements software.

### Transmission EM

Vero E6-hAT cells were seeded at 0.5 x 10^6^ cells per well in permanox dishes, infected with WT virus (MOI=0.1) or ΔE virus (MOI=0.3), or mock infected. After 12 h, cell monolayers were rinsed once with PBS, followed by fixation *in situ* with 2.5% glutaraldehyde in PBS (pH 7.4) for 1 h at room temperature. Samples were then washed three times in PBS for 10 min each. Post-fixation was carried out in 1% osmium tetroxide (OsO ) containing 1% potassium ferricyanide for 1 h at 4° C. Monolayers were again washed three times in PBS for 10 min each. Samples were dehydrated through a graded ethanol series consisting of 30%, 50%, 70%, and 90% ethanol (10 min each), followed by three changes in 100% ethanol for 15 min each. Dehydrated monolayers were infiltrated with epoxy resin (Epon) through three sequential 1 h exchanges. After the final resin change, excess resin was removed, and embedding was performed by inverting resin-filled beam capsules directly onto selected regions of the monolayer. Polymerization was carried out overnight at 37° C and subsequently for 48 h at 60° C. Following polymerization, the cured beam capsules along with the attached monolayer were separated from the permanox dish surface. Blocks were trimmed and ultrathin-sectioned using a Leica Ultracut R ultramicrotome, and images were acquired on a JEM-1400 transmission electron microscope.

### Statistics

Unless otherwise specified, results were analyzed for statistical significance by two-sided Welch’s t test with Prism software (GraphPad). A p-value of less than or equal to 0.05 was used to indicate statistical significance.

## Acknowledgements

This work was supported by National Institutes of Health (NIH) grants HL129949 (H.L.F), AI180311 (W.P.D.), AI175795 (Z.A.), and the Pittsburgh CTSI COVID-19 Pilot Program (Z.A.). The Pittsburgh Center for CryoEM (RRID:SCR_025216) used for data collection in this project was supported, in part, by the University of Pittsburgh, the School of Medicine, and the Department of Structural Biology.

