## Supplemental figures and legends for "Deletion of the Envelope gene attenuates SARS-CoV-2 infection by altered Spike localization and increased cell-to-cell transmission"

#### Supplemental Figure Legends

**Figure S1. Design of a BAC-based SARS-CoV-2 virus production system.** (A) Schematic of the assembly of the SARS-CoV-2 BAC. (B) Workflow for production of SARS-CoV-2 virus via BAC transfection.

**Figure S2**. **WT and ΔE SARS-CoV-2 viruses lacking the nLuc reporter gene produce similar results to WT and ΔE SARS-CoV-2-nLuc viruses.** (A) Vero E6-hAT cells were infected with WT SARS-CoV-2 virus (MOI=0.01) or mock infected for 24 h and imaged by phase contrast microscopy with a 10X objective. Solid white bar represents 100 μm. (B) Plaque assays are shown for WT SARS-CoV-2 infection of Vero E6-hAT cells and ΔE SARS-CoV-2 infection of Vero E6-hAT cells with or without stable expression of E. Numbers represent log dilution of virus. (C-D) Vero E6-hAT cells (WT and ΔE) or E6-hAT cells with stable expression of E (ΔE only) were infected with WT or ΔE SARS-CoV-2 virus for 72 h. Plaque titers (C) and genomic RNA (ORF1a) copies (D) were measured. Dotted lines represent the assay limits of detection. * $p < 0.05$, ns $p > 0.05$. Error bars represent SEM for two independent experiments. (E) Vero E6-hAT cells with or without stable expression of E were infected with WT or ΔE SARS-CoV-2 (MOI=0.01) for 24 h, stained for S, and imaged by widefield epifluorescence microscopy. Solid white bars represent 100 μm.

**Figure S3. Production and assay of ΔE SARS-CoV-2 virus.** (A) Schematic of E deletion from the SARS-CoV-2 genome. (B) coE was cloned downstream from the T2A peptide and codon-optimized mRuby3 in a doxycycline-inducible lentiviral vector. Widefield epifluorescence microscopy was used to visualize mRuby3 expression in stably transduced cells with and without doxycycline induction. Solid white bars represent 40 μm. (C) Immunoblot of coE expression in BHK and Vero E6-hAT cells stably expressing coE with and without doxycycline induction. (D) Workflow for production of ΔE SARS-CoV-2 virus via BAC transfection with and without coE trans-complementation. (E) Workflow for passaging and characterizing ΔE SARS-CoV-2 virus.

**Figure S4.** RNA quantitation of endogenous E (A) or trans-complemented coE (B) copies were measured by qRT-PCR in producer cells (left) and virus (right) from serial passages of WT and ΔE SARS-CoV-2-nLuc shown in **Figure 2**. Dotted lines represent the assay limits of detection. Error bars represent SEM for three independent experiments.

**Figure S5.** Partial rescue of ΔE SARS-CoV-2 infectivity attenuation requires E expression in infected cells. (A) Vero E6-hAT cells were infected with WT or ΔE SARS-CoV-2-nLuc produced from cells with or without E trans-complementation and assayed for nLuc activity after 72 h. Error bars represent SEM for three independent experiments. One-way ANOVA with Tukey's multiple comparison's test was used to determine statistical significance. ns $p > 0.05$. (B) Vero E6-hAT cells with or without stable expression of E were infected with WT or ΔE SARS-CoV-2-nLuc and assayed for nLuc activity after 24 h. Error bars represent SEM for two independent experiments. ns $p > 0.05$. Dotted lines represent the assay limits of detection.

**Figure S6**. Detection of syncytia induced by SARS-CoV-2 infection. (A) Schematic and workflow for visualizing syncytia via the Split-eGFP BiFC reporter cell system. (B) Vero E6-hAT-eGFP-N and Vero E6-hAT-eGFP-C BiFC reporter cells co-cultured 1:1 with and without stable expression of E were infected with WT or ΔE SARS-CoV-2-nLuc virus for 1 h, treated with Nab for 24 h, stained for nLuc expression (red) and Hoechst (blue), and imaged by confocal microscopy. Reconstituted eGFP is shown in green. mRuby3 expression was also assessed (white). Representative micrographs from two independent experiments are shown. Solid white bars represent 50 μm.

**Figure S7.** Lack of E during SARS-CoV-2 infection results in ERGIC reorganization. Vero E6-hAT cells with stable E expression were infected with WT or ΔE SARS-CoV-2-nLuc for 24 h, stained for ERGIC-53 (green), S (red) and Hoechst (blue), and imaged by confocal microscopy. mRuby3 expression was also assessed (white). Solid white bars represent 20 μm.

### Figure S1

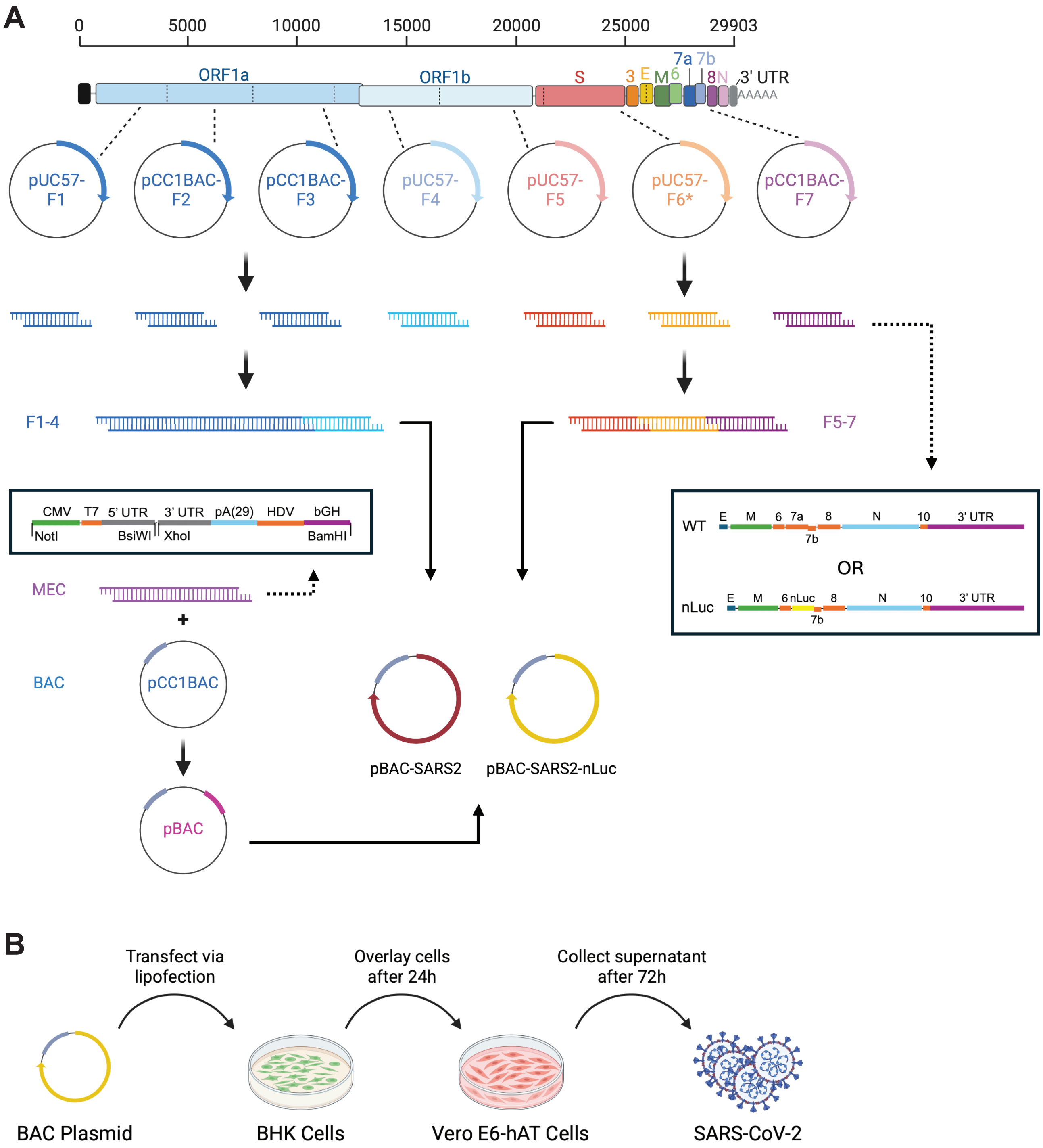
A
0
5000
10000
15000
20000
25000
29903
ORF1a
ORF1b
S
3
E
M
6
7a
7b
8
N
3' UTR
AAAAA
pUC57-F1
pCC1BAC-F2
pCC1BAC-F3
pUC57-F4
pUC57-F5
pUC57-F6*
pCC1BAC-F7
F1-4
F5-7
CMV
T7
5' UTR
3' UTR
pA(29)
HDV
bGH
NotI
BsiWI
XhoI
BamHI
MEC
+
BAC
pCC1BAC
pBAC
pBAC-SARS2
pBAC-SARS2-nLuc
WT
E
M
6
7a
8
N
10
3' UTR
7b
OR
nLuc
E
M
6
nLuc
8
N
10
3' UTR
7b
B
Transfect via lipofection
Overlay cells after 24h
Collect supernatant after 72h
BAC Plasmid
BHK Cells
Vero E6-hAT Cells
SARS-CoV-2

### Figure S2

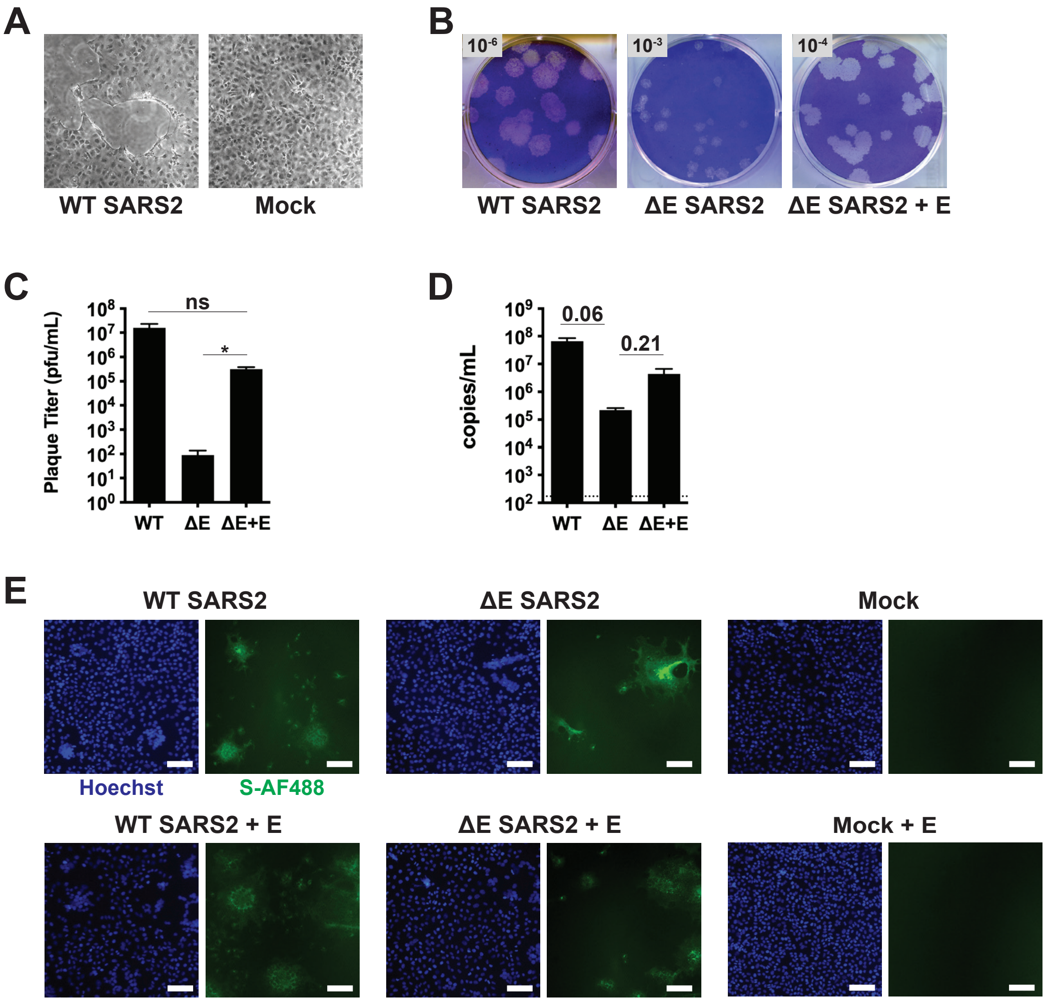

A
WT SARS2
Mock
B
10-6
10-3
10-4
WT SARS2
ΔE SARS2
ΔE SARS2 + E
C
Plaque Titer (pfu/mL)
ns
*
WT
ΔE
ΔE+E
D
copies/mL
0.06
0.21
WT
ΔE
ΔE+E
E
WT SARS2
ΔE SARS2
Mock
Hoechst
S-AF488
WT SARS2 + E
ΔE SARS2 + E
Mock + E

### Figure S3

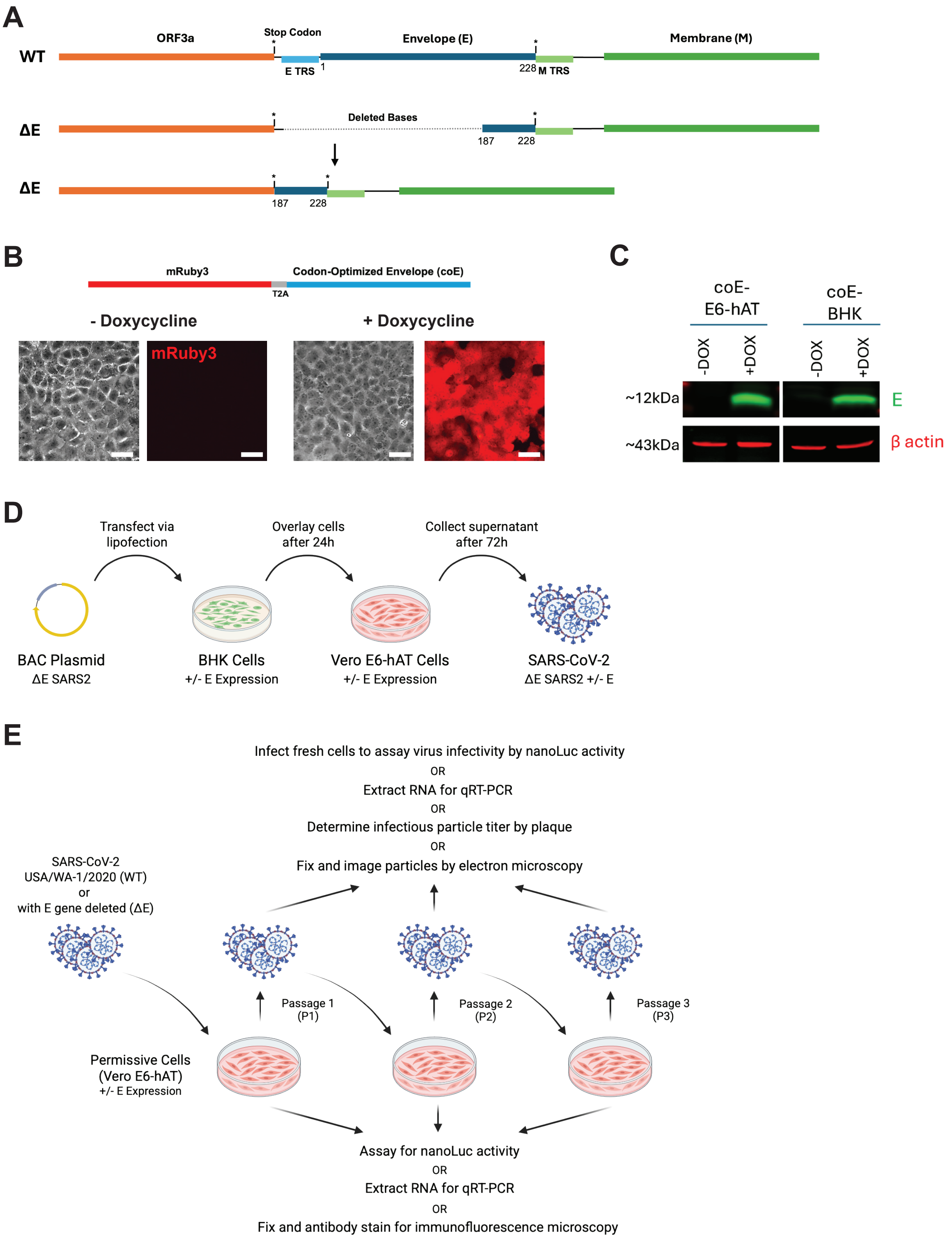
A
WT
ORF3a
Stop Codon
Envelope (E)
Membrane (M)
E TRS
1
228
M TRS
ΔE
Deleted Bases
187
228
ΔE
187
228
B
mRuby3
Codon-Optimized Envelope (coE)
T2A
- Doxycycline
+ Doxycycline
mRuby3
C
coE-
E6-hAT
coE-
BHK
-DOX
+DOX
-DOX
+DOX
~12kDa
E
~43kDa
β actin
D
Transfect via
lipofection
Overlay cells
after 24h
Collect supernatant
after 72h
BAC Plasmid
ΔE SARS2
BHK Cells
+/- E Expression
Vero E6-hAT Cells
+/- E Expression
SARS-CoV-2
ΔE SARS2 +/- E
E
Infect fresh cells to assay virus infectivity by nanoLuc activity
OR
Extract RNA for qRT-PCR
OR
Determine infectious particle titer by plaque
OR
Fix and image particles by electron microscopy
SARS-CoV-2
USA/WA-1/2020 (WT)
or
with E gene deleted (ΔE)
Passage 1
(P1)
Passage 2
(P2)
Passage 3
(P3)
Permissive Cells
(Vero E6-hAT)
+/- E Expression
Assay for nanoLuc activity
OR
Extract RNA for qRT-PCR
OR
Fix and antibody stain for immunofluorescence microscopy

### Figure S4

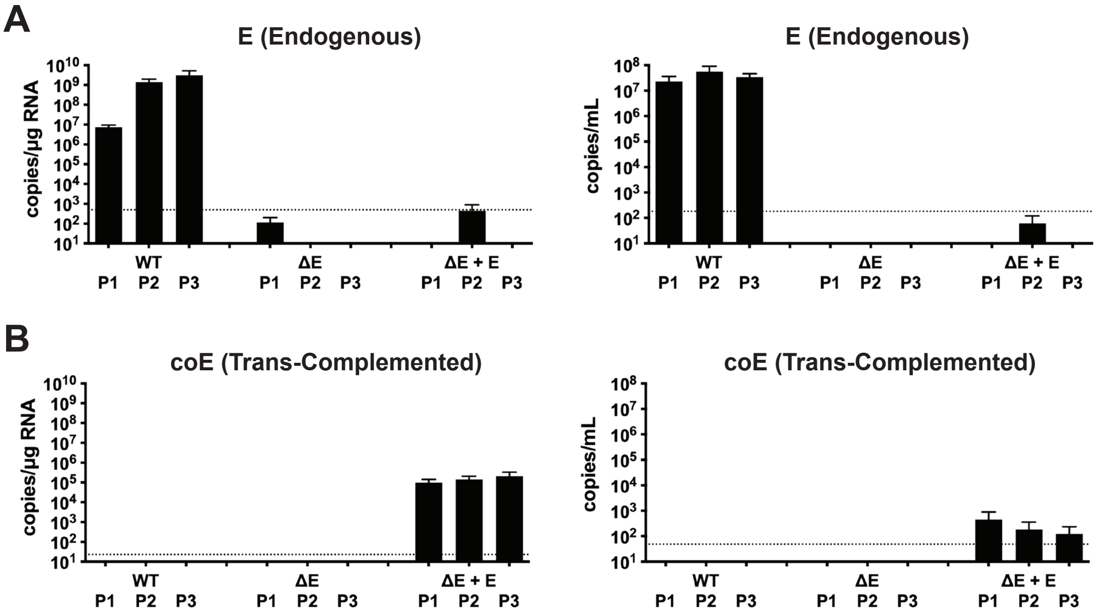

A
E (Endogenous)
copies/µg RNA
copies/mL
WT
ΔE
ΔE + E
P1 P2 P3 P1 P2 P3 P1 P2 P3
B
coE (Trans-Complemented)
copies/µg RNA
copies/mL
WT
ΔE
ΔE + E
P1 P2 P3 P1 P2 P3 P1 P2 P3

### Figure S5

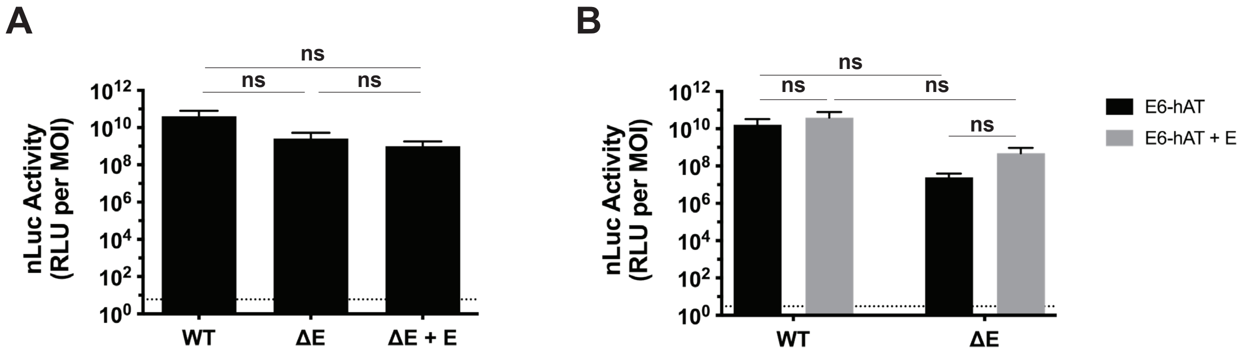
A
B
ns
ns
ns
nLuc Activity
(RLU per MOI)
$10^{12}$
$10^{10}$
$10^{8}$
$10^{6}$
$10^{4}$
$10^{2}$
$10^{0}$
WT
ΔE
ΔE + E
E6-hAT
E6-hAT + E

### Figure S6

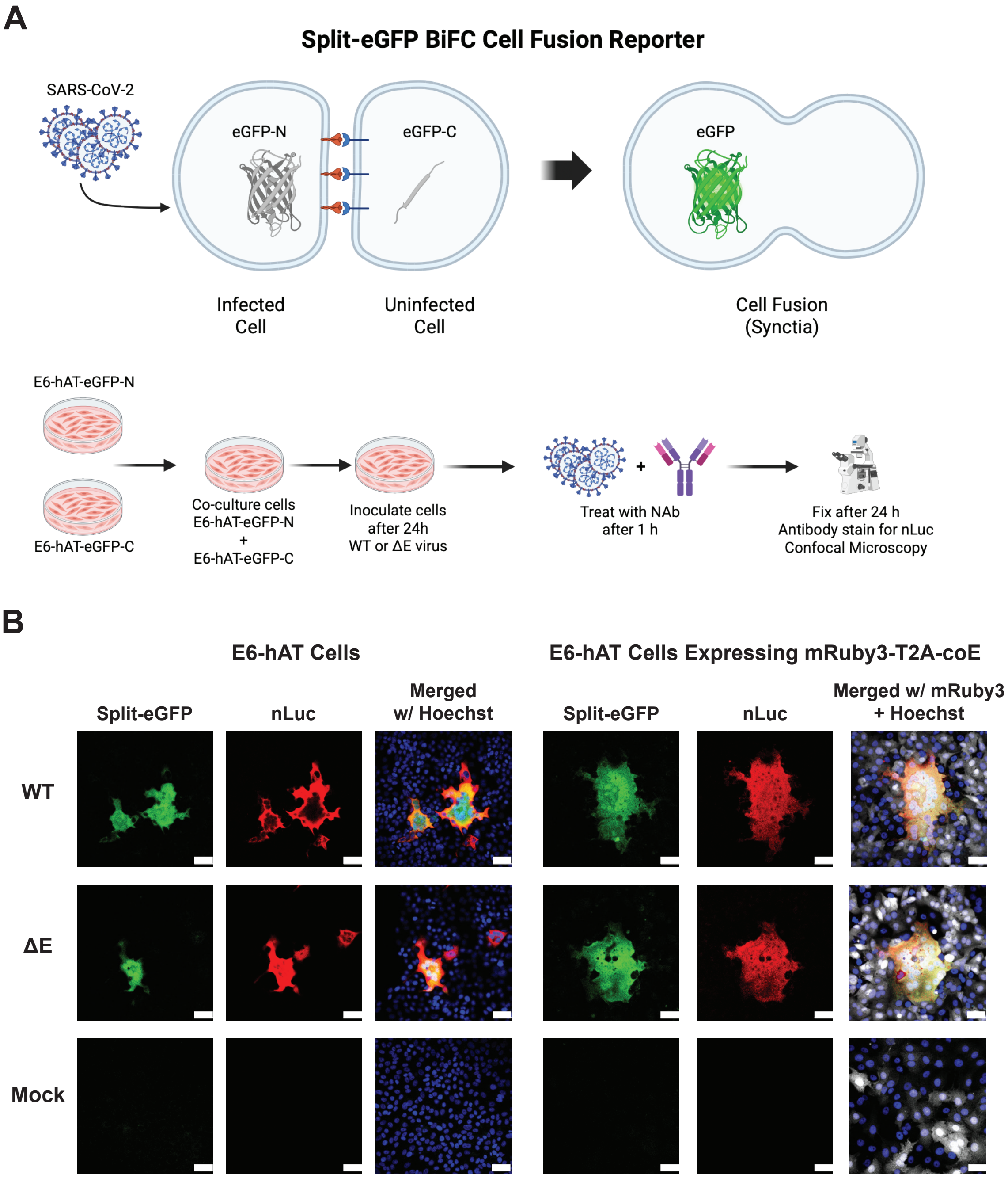
A
Split-eGFP BiFC Cell Fusion Reporter
SARS-CoV-2
eGFP-N
eGFP-C
eGFP
Infected Cell
Uninfected Cell
Cell Fusion (Syncytia)
E6-hAT-eGFP-N
E6-hAT-eGFP-C
Co-culture cells E6-hAT-eGFP-N + E6-hAT-eGFP-C
Inoculate cells after 24h WT or ΔE virus
+
Treat with NAb after 1 h
Fix after 24 h Antibody stain for nLuc Confocal Microscopy
B
E6-hAT Cells
E6-hAT Cells Expressing mRuby3-T2A-coE
Split-eGFP
nLuc
Merged w/ Hoechst
Split-eGFP
nLuc
Merged w/ mRuby3 + Hoechst
WT
ΔE
Mock

### Figure S7

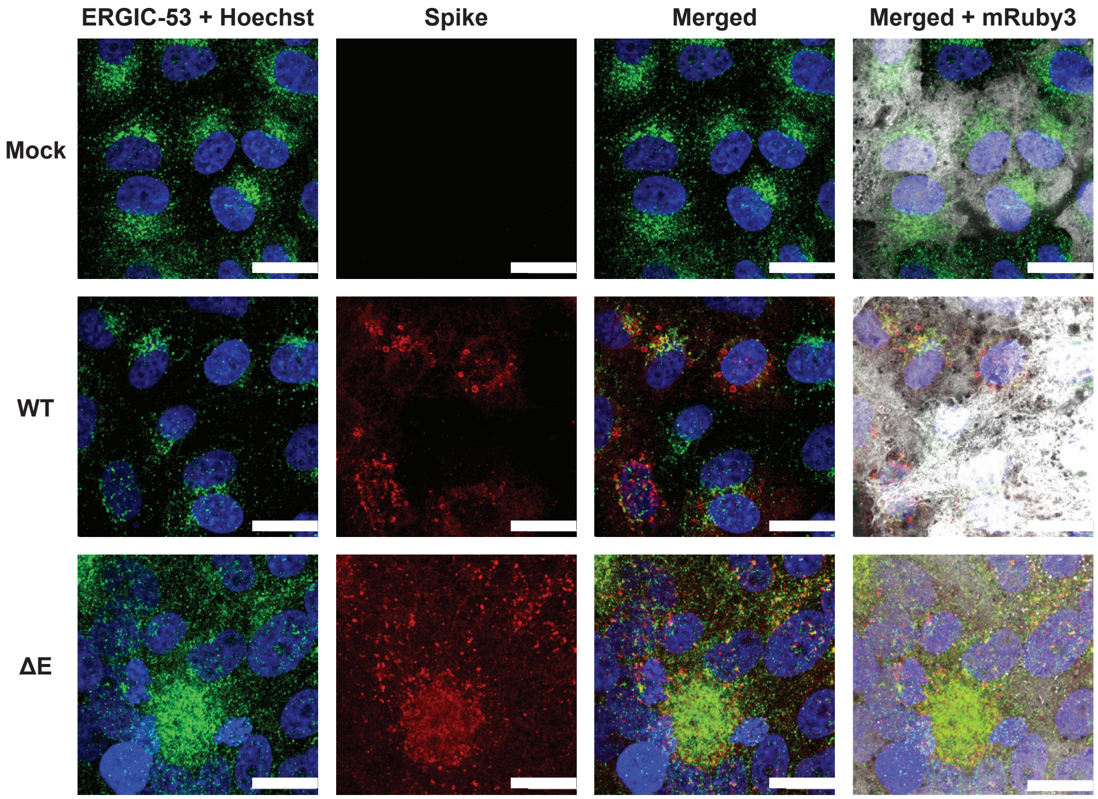
